# Melanoblast transcriptome analysis reveals novel pathways promoting melanoma metastasis

**DOI:** 10.1101/721712

**Authors:** Kerrie L. Marie, Antonella Sassano, Howard H. Yang, Aleksandra M. Michalowski, Helen T. Michael, Theresa Guo, Yien Che Tsai, Allan M. Weissman, Maxwell P. Lee, Lisa M. Jenkins, M. Raza Zaidi, Eva Pérez-Guijarro, Chi-Ping Day, Heinz Arnheiter, Sean Davis, Paul S. Meltzer, Glenn Merlino, Pravin J. Mishra

## Abstract

Cutaneous malignant melanoma is an aggressive cancer of melanocytes with a strong propensity to metastasize. We posited that melanoma cells acquire metastatic capability by adopting an embryonic-like phenotype, and that a lineage approach would uncover novel metastatic melanoma biology. We used a genetically engineered mouse model to generate a rich melanoblast transcriptome dataset, identified melanoblast-specific genes whose expression contributed to metastatic competence, and derived a 43-gene signature that predicted patient survival. We identified a melanoblast gene, *KDELR3*, whose loss impaired experimental metastasis. In contrast, *KDELR1* deficiency enhanced metastasis, providing the first example of different disease etiologies within the KDELR-family of retrograde transporters. We show that KDELR3 regulates the metastasis suppressor, KAI1, and report an interaction with the E3 ubiquitin-protein ligase gp78, a regulator of KAI1 degradation. Our work demonstrates that the melanoblast transcriptome can be mined to uncover novel targetable pathways for melanoma therapy.

---

Melanoma is an aggressive cancer that frequently progresses to metastatic proficiency. Treatment of metastatic melanoma remains a challenge, highlighting an urgent need to uncover new targets that could be used in the clinic to broaden therapeutic options. In the early 19^th^ century, Virchow first described cancer cells as being “embryonic-like”^1^. Developmental systems have since proven useful to study melanoma, and melanoma cell plasticity appears to be a key feature of melanoma progression. Melanocyte lineage pathways are a recurring theme in melanoma etiology, reinforcing the importance of uncovering new melanocyte developmental pathways and biology^2-13^. Here we use a genetically engineered mouse (GEM), designed to facilitate the isolation and analysis of developing melanocytes (melanoblasts), to attempt to uncover new targets relevant to melanoma metastasis.

Melanocytes are neural crest-derived cells whose development necessitates extensive migration/invasion to populate the skin and other sites^14^. This process requires melanoblasts to adopt a migratory phenotype, to interact with and survive in foreign microenvironments and to colonize distant sites − functions that are analogous to metastatic competence^15^. To complete these processes, the cell may encounter numerous cellular stressors, such as shear stress, nutrient deprivation, hypoxia, lipid stress and oxidative stress^16^. The cellular impact of these stressors converges at the Endoplasmic Reticulum (ER), the organelle tied closely to protein synthesis and responsible for correct protein folding, protein quality control and post-translational modifications. Stress stimuli can result in aberrant ER function, a build-up of unfolded/misfolded proteins (ER stress), and an overwhelmed system. The ER can therefore be viewed as an exquisitely sensitive stress sensor. Upon ER stress insult, the ER launches an immediate counter measure known as of the Unfolded Protein Response (UPR)^17^. The UPR consists of three arms, the IRE1, PERK and ATF6 pathways. Cumulatively these result in transcriptional activation of chaperones and ER-Associated Degradation (ERAD) machinery that target unfolded proteins for degradation to help counter the stress^17^. Simultaneously the PERK pathway initiates general translation attenuation to reduce protein load in the ER. Un-checked levels of ER stress result in cell death via the PERK-stimulated CHOP pathway^17^. The KDEL-Receptors (KDELRs) are a family of seven-transmembrane-domain ER protein retention receptors consisting of three members (KDELR1, 2 and 3) that function in the ER Stress Response (ERSR). They share structural homology, but each isoform can have different ligands^18, 19^. They are responsible for the retrograde transport of protein machinery from the Golgi to the ER, including chaperones that target unfolded proteins for re-folding, and whose disassociation from membrane receptors stimulates UPR signaling^19, 20^. In embryogenesis, there is a need for tightly coordinated temporal control of gene/protein expression for correct differentiation of tissues^16^. Embryonic cells are therefore primed to accommodate overwhelming ER stress, as this would affect the cell’s ability to translate, synthesize, fold and modify proteins, which would compromise the developing embryo^16^.

We hypothesized that genes whose expression is upregulated in developing melanoblasts and metastatic melanoma but downregulated in differentiated melanocytes (hereafter referred to as MetDev genes), can be reactivated by melanoma cells to facilitate metastasis (Fig. 1a). To explore this, we took advantage of a GEM model in which GFP is inducibly targeted to embryonic melanoblasts and mature melanocytes by using the Dopachrome tautomerase (*Dct*) promoter to drive expression (inducible *Dct*-GFP; i*Dct*-GFP)^21, 22^. This powerful tool enables identification and isolation of cells of the melanocytic lineage^21^, which can be employed to investigate the melanoblast transcriptome. Using this approach, we identified a 43-gene embryonic melanoblast gene signature that predicts metastatic melanoma patient survival, and we highlight a new role for KDELR3^20^, distinct from other members of the KDELR family. A metastasis suppressor screen highlights KAI1/CD82 (hereafter referred to as KAI1) as a KDELR3-regulated protein. We observe that KDELR3 regulates KAI1 protein levels and post-translational modification. We demonstrate an undescribed interaction of KDELR3 with gp78, the E3 ubiquitin protein ligase known to regulate KAI1 degradation^23^. Our work shows that melanoma cells can commandeer embryonic transcriptomic programs to promote their progression to metastasis. These genes represent an untapped source of novel targetable pathways to exploit for improving melanoma treatment.

**Figure 1:**
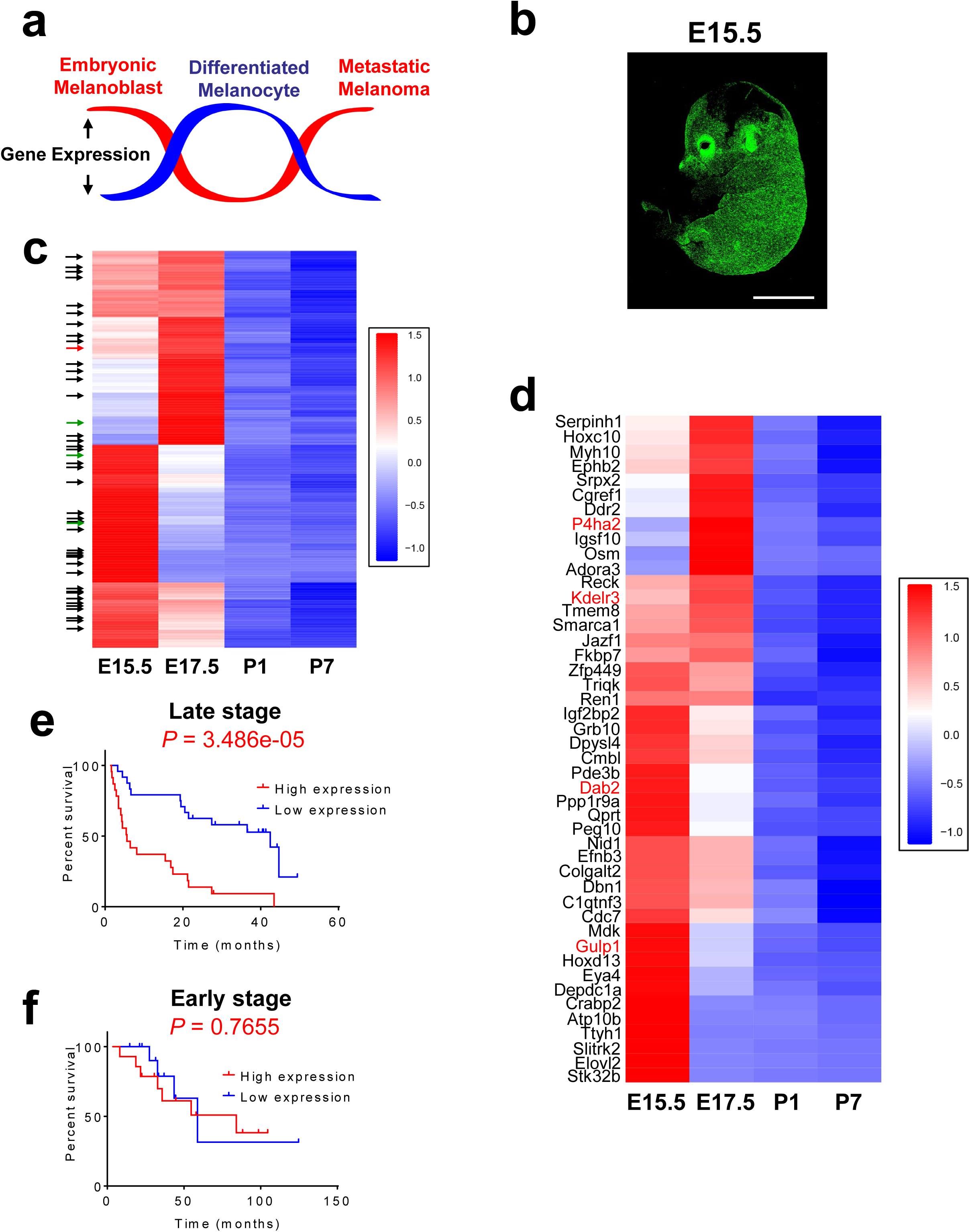
Discovery of metastasis-development (MetDev) genes. **a**, Schematic depicting the experimental hypothesis: genes whose expression is upregulated in melanoblasts and metastatic melanoma, but downregulated in differentiated melanocytes (red line), may drive cellular functions that promote melanoma metastasis (MetDev genes). **b**, Confocal imaging of *i*Dct-GFP embryo, Embryonic day 15.5 is x5, scale bars, 5 mm. **c**, RNA-Seq expression of mouse developing melanocytes: 467 embryo-specific genes shown. Black arrows: 43 genes identified from Cox proportional hazards model. Green arrows: Genes functionally validated. Red arrow: *Kdelr3*. Embryonic day 15.5 and 17.5 (E15.5 and E17.5 respectively). Postnatal day 1 and postnatal day 7 (P1, P7 respectively). **d**, RNA-Seq expression of 46 genes in mouse developing melanocytes: Black text: 43 genes identified from Cox proportional hazards model. Red text: 4 genes functionally validated. *Kdelr3* validated both in Cox proportional hazards model and functionally validated. **e**-**f**, Cox proportional hazards modeling (GSE19234) yielded a 43-gene MetDev signature. Patients’ risk assessed in GSE8401 patient cohort. Late: stage III/IV metastatic melanomas. Early: stage I/II primary tumors. High: high expression of gene signature. Low: low expression of gene signature. Log rank test. Late stage, high (N = 23) vs. low (N = 24), *P* = 3.486e-05. Early stage, high (N = 14) vs low (N = 13), *P* =0.7655.

## Results

### Melanoblast transcriptomic expression in melanoma metastasis

To study melanoblast genes, GFP-positive melanocytic cells were isolated from four developmental time points: Embryonic day (E) 15.5 and 17.5, and Postnatal day (P) 1 and 7 (Fig. 1b, Supplementary Fig. 1a-b). These four stages represent embryonic melanoblast development from the neural crest into differentiated quiescent melanocytes of the postnatal pup^24, 25^. Melanocytes/ melanoblasts were isolated using Fluorescence-Activated Cell Sorting (FACS) from i*Dct*-GFP mice (Supplementary Fig. 1c). At E15.5 and E17.5 melanoblasts are still migrating and colonizing the hair follicles within the epidermis^24-26^ − processes that we believe are highly relevant to metastasis, particularly with respect to colonization at the metastatic site – and intrafollicular melanoblasts are still present^26^. P1 and P7 mature melanocytes were selected as a model of quiescent differentiated melanocytes; these time points are prior to the first hair follicle cycle that begins at 6 weeks post birth. Melanocytic cells were extracted from multiple litters (6-10 pups) at each developmental stage to ensure comprehensive representation of all melanoblasts/ melanocytes present. RNA was extracted for whole-transcriptome sequencing.

Genes with differential expression between embryonic melanoblasts (E15.5 and E17.5) and post-natal differentiated melanocytes (P1 and P7) were identified using DESeq2^27^ with a q-value < 0.1, yielding 976 differentially expressed genes (Supplementary Fig. 2). Of these genes, we filtered out any whose differential expression was less than 1.5 log2 fold increased in melanoblasts, as we deemed that a fold change of less than this was unlikely to be biologically meaningful. 467 melanoblast-specific genes were identified from our analyses, which we hypothesize to be putative melanoma metastasis enhancer genes (MetDev genes; Fig. 1c). To test the relevance of our melanoblast gene cohort in melanoma metastasis we interrogated this gene list in melanoma patient data. To ask if our 467-gene MetDev cohort was enriched in genes that contributed to poor progression of patients, we used a Cox proportional hazards model to associate their expression with overall survival in a training dataset of human patient samples derived from melanoma metastases (stage III and stage IV; GSE19234)^28^. We discerned a 43-gene survival risk predictor (Fig. 1c, black/red arrows; Fig. 1d, black text, *Kdelr3*) that could accurately predict patient outcome in a separate testing dataset of late stage (stage III and stage IV) metastatic melanoma patient samples derived from metastases (GSE8401; Fig. 1e)^29^. These data show that not only is our MetDev cohort enriched for metastatic progression genes, but it can also predict survival in multiple independent patient datasets. Notably, gene expression levels in samples derived from early stage (stage I and stage II) primary melanoma lesions did not predict patient outcome, suggesting that MetDev genes play a key role in late-stage disease specifically (GSE8401; Fig. 1f)^29^.

To allow functional validation of our MetDev candidates in both soft agar colony forming assays and in experimental metastasis models we elected to prioritize the list of MetDev gene candidates. To do this in an unbiased fashion we applied criteria based solely on melanoblast expression data, selecting for genes with no detectable gene expression in P7 postnatal pups. Differential expression was validated using a separate microarray expression dataset derived from our i*Dct*-GFP model (E17.5 vs P2 and P7; q-value < 0.1)^21^. Further criteria using differences in fold-increase expression in melanoblasts vs. melanocytes and the greatest expression at embryonic stages allowed us to select 20 genes likely to be the most functionally relevant. Of these 20 we noted that 7 genes (*Kdelr3*, *P4ha2*, *Gulp1*, *Dab2*, *Lum*, *Aspn*, *Mfap5*) were associated with Extracellular Matrix (ECM) or trafficking. For functional analyses, we chose 4 of these 7 genes (*Kdelr3*, *P4ha2*, *Gulp1, Dab2*) with no established role in cutaneous melanoma metastasis (Fig. 1c, green/red arrows; Fig. 1d, red text). siRNA knockdown of our four candidate genes in B16 mouse melanoma cells inhibited both growth in soft agar colony formation assays and formation of lung metastases in experimental metastasis assays compared to non-targeting controls (Table 1). Our work demonstrates that the MetDev dataset is enriched in genes that have a functional role in melanoma metastasis. We identify four new melanoma metastasis genes and highlight ECM and trafficking as important pathways common to both melanoblast development and melanoma metastasis.

**Table 1:** siRNA screen for metastatic potential of four putative MetDev genes. siRNA knockdown of genes indicated (B16 cell line). Colony formation assay, n = 10 wells (*Dab2*, *Kdelr3*, Control), n = 5 wells (*Gulp1*, *P4ha2*, Control), screen performed once. *P*-value assessed by Kruskal-Wallis using uncorrected Dunn’s test versus siControl. Tail vein metastasis assay, n = 10 mice (*Dab2*, *Kdelr3*, Control), n = 5 wells (*Gulp1*, *P4ha2*, Control), screen performed once. *P*-value assessed by Kruskal-Wallis using uncorrected Dunn’s test versus siControl.

| Gene Symbol | Colony Formation | Tail vein metastasis |
| --- | --- | --- |
| Gulp1 | P = 0.0002 | P = 0.0002 |
| Kdelr3 | P = 0.0122 | P = 0.0155 |
| P4ha2 | P = 0.022 | P = 0.022 |
| Dab2 | P = 0.0428 | P = 0.0421 |

We further observed significant co-expression of three of the four functionally validated genes (*Kdelr3, P4ha2* and *Dab2*) throughout four distinct mouse models of melanoma (See Methods and Supplementary Table 1), corroborated in a melanoma patient cohort (TCGA; Supplementary Table 2). Notably, expression of *Kdelr3* and *P4ha2* was highly correlated throughout all datasets (Supplementary Fig. 3a-b), raising the possibility that some metastasis-associated MetDev genes may be co-regulated and serve a more coordinated role in metastasis.

### *KDELR3* is a Golgi-resident protein whose expression correlates with melanoblast development and melanoma progression

To understand how melanoblast genes might facilitate metastasis we chose to study one MetDev gene in depth. *KDELR3* was selected as it was a positive hit in all of our analyses: *KDELR3* is a trafficking protein important in the ERSR whose expression was associated with poor patient prognosis in metastatic melanomas (Fig. 1e, 43 gene signature), and *KDELR3* was functionally validated in both soft agar colony formation and experimental metastasis assays (Table 1). The KDELRs are Golgi-to-ER retrograde transporters responsible for maintaining ER localization of their protein substrates. KDELR substrates consist of protein chaperones required for protein folding and targeting unfolded proteins for degradation^20^, thereby assisting the UPR and maintaining ER quality in times of ER stress. We show that KDELR3 is localized to both the cis- and trans-Golgi compartments in metastatic melanoma cells (Supplementary Fig. 3c) and validate expression of KDELR3 in mouse melanoblasts (Fig. 2a). Moreover, within the KDELR family only *KDELR3* demonstrated a melanoblast-specific expression pattern and showed consistent upregulation in melanoma cell lines (Fig. 2b; Supplementary Fig. 3d-e). These data raise the possibility that *KDELR3* plays a role in melanoma progression which is distinct from other KDELRs, despite their presumed redundancy. Analysis of human patient datasets and tumor histology microarrays confirmed an upregulation of *KDELR3* expression in malignant melanoma vs. benign nevi (Fig. 2c-e).

**Figure 2:**
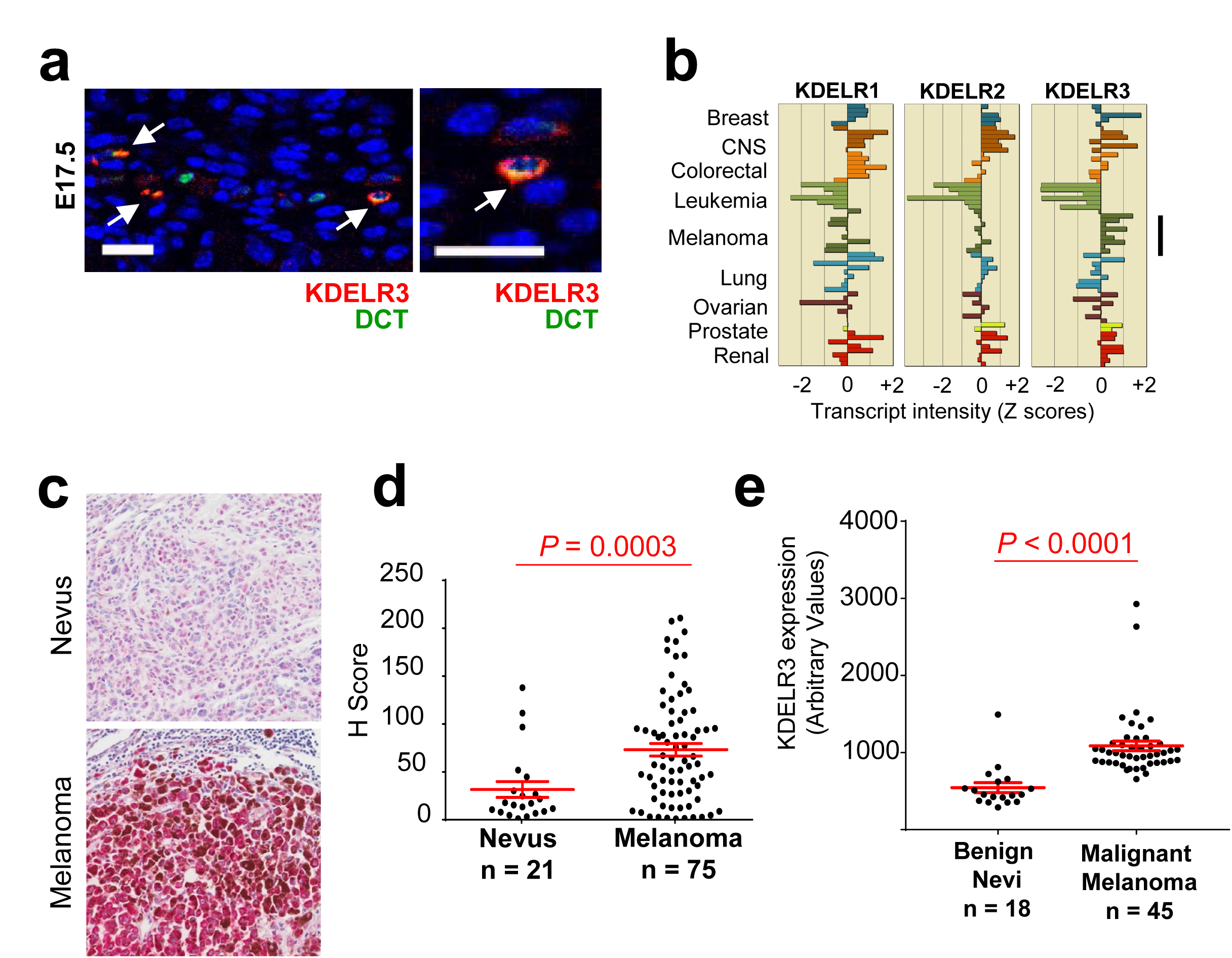
Melanoblast gene expression in melanoma. **a**, KDELR3 (red) and DCT (green) staining in E17.5 mouse skin. White arrows depict co-localization. Magnification, 40x. Scale bars, 20µm. Representative image of 100 cells analyzed taken from one mouse. **b**, Pan-cancer RNA expression of KDEL Receptors in human cell lines (NCI60; CellMiner analysis); *KDELR3* expression in melanoma (black line). **c**, *KDELR3* expression in human nevus and melanoma lymph node metastasis (red intercellular staining), magnification, 20x. **d**, H-Score of KDELR3 immunohistochemistry in human tumor microarrays. Unpaired two-tailed student’s t-test with Welch’s correction, *P* = 0.0003, *df* = 47.9, *t* = 3.936. **e**, *KDELR3* expression in benign nevi and malignant melanoma (GSE3189; 204017_at probeset). Unpaired two-tailed student’s t-test with Welch’s correction, *P* < 0.0001, *df* = 47.39, *t* = 6.035. **d**, **e**, Line and error bars represent mean ± s.e.m.

We sought to functionally validate a role for *KDELR3* in melanoma progression. We used human and mouse melanoma cells to demonstrate that small-interfering RNA (siRNA) and short-hairpin RNA (shRNA) knockdown of *KDELR3* significantly reduced, and *KDELR3* overexpression enhanced, anchorage-independent growth (Fig. 3a-d; Supplementary Fig. 4a-b), which cannot be attributed to a change in proliferation (Supplementary Fig. 4c). There are two *KDELR3* variants, and we selected the *KDELR3-001* variant to perform rescue experiments as it is the most abundant transcript expressed in human cell lines and patient samples. We therefore performed rescue experiments via exogenous expression of *KDELR3-001^Mu^*, whose shRNA recognition site had been mutated without altering the final protein sequence. *KDELR3-001^Mu^* expression was restored, rescuing the anchorage-independent growth phenotype (Fig. 3e-g; Supplementary Fig. 4d). *KDELR3* was therefore validated as a mediator of anchorage-independent growth in melanoma cells, a process required for metastasis.

**Figure 3:**
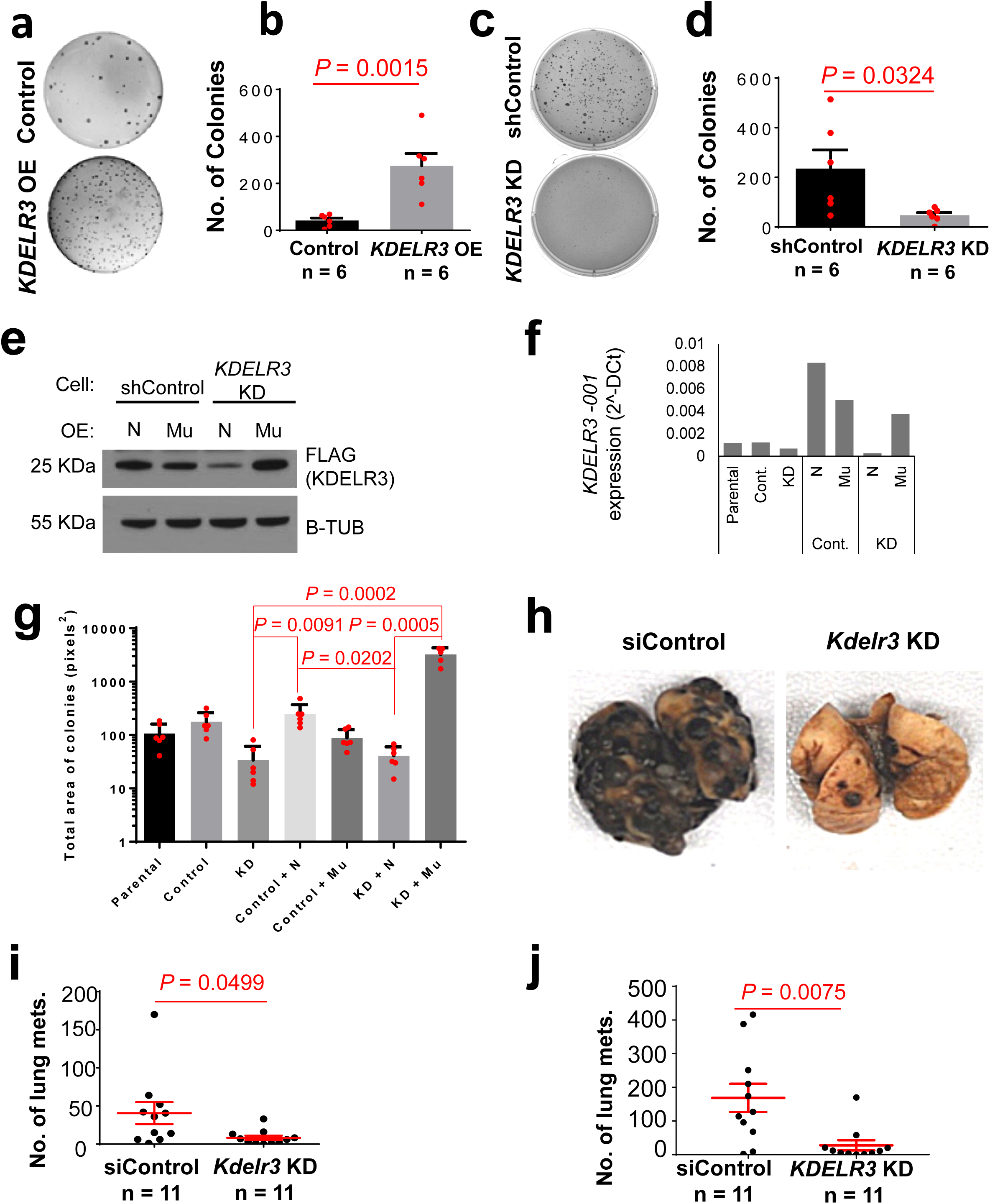
*KDELR3* mediates melanoma metastatic potential. **a-d**, Soft agar colony formation assay with: **a**-**b**, overexpression of *KDELR3* in human SK-MEL-28 cells versus parental cell, unpaired two-tailed student’s t-test, *P* = 0.0015, *df* = 10, *t* =4.307. 6 wells analyzed per group. **c**-**d**, shRNA *KDELR3* knockdown in human WM-46 cells versus non-targeting control, unpaired two-tailed student’s t-test, *P* = 0.0324, *df* = 10, *t* =2.483. 6 wells analyzed per group. **e**-**f**, Western blot and qPCR analysis of exogenously expressed FLAG-tagged *KDELR3-001*; ENST00000216014 (N) and *KDELR3-001^Mu^* (Mu) in WM-46 (**e**) and 1205Lu (**f**) cells, transduced with non-targeting control (shControl/Cont./Control) or *KDELR3*-targeted (KD) shRNAs. Total *KDELR3-001* RNA (*KDELR3-001* and *KDELR3-001^Mu^*) (**f**). **g**, Rescue of soft agar colony formation in *KDELR3-001^Mu^* cells (WM-46), Kruskal-Wallis with Dunn’s multiple comparison test. 5-6 wells analyzed per group. **h**-**i**, Tail vein metastasis of *Kdelr3* siRNA knockdown in mouse B16 cells. Unpaired two-tailed student’s t-test with Welch’s correction, *P* =0.0499, *df* = 10.83, *t* = 2.207. **j**, Tail vein metastasis of *KDELR3* siRNA-mediated knockdown human 1205Lu cells transduced with *Ferh-luc-GFP*. Unpaired two-tailed student’s t-test with Welch’s correction, *P* = 0.0075, *df* = 12.57, *t* = 3. **b**, **d**, **e**, **i**, **j**, Bars and error bars depict mean + s.e.m. **a**-**f**, **h**-**j**, Representative of three independent experiments. **g**, Representative of two independent experiments. **e**, β-Tubulin loading control.

### *KDELR3* knockdown reduces lung colonization in experimental metastasis assays

To assess the relevance of *KDELR3* within the metastatic cascade, we used a tail vein experimental metastasis assay, which specifically assesses the ability of the cells to extravasate and colonize the lung, processes that are critical for metastatic capacity. Tail vein metastasis assays enable lung colonization to be assessed with greater specificity/sensitivity − biology that we suggest may be mirrored during hair follicle colonization (E17.5). Transient knockdown of *KDELR3* in either mouse (Fig. 3h-i) or human melanoma cell lines (Fig. 3j, Supplementary Fig. 5a) resulted in significantly reduced metastatic potential compared to non-targeting controls, indicating that *KDELR3* expression is important for the cells’ ability to extravasate/ colonize the lung, further validating that *KDELR3* is a melanoblast gene that functions in metastasis (MetDev gene). Stable shRNA knockdown of *KDELR3* also resulted in a reduction in lung colonization following tail vein metastasis and significantly fewer mice characterized with high metastatic burden (Supplementary Fig. 5b-f). However, no appreciable difference in cell cycle or subcutaneous *in vivo* tumor growth was observed (Supplementary Fig. 5g-i), suggesting that the *KDELR3*-mediated metastatic phenotype cannot be attributed to a change in proliferation, and that *KDELR3* is a genuine melanoma metastasis progression gene.

### *KDELR3* and the ER Stress Response in metastatic melanoma

To uncover how *KDELR3* expression may be involved with melanoma metastasis, we asked which pathways were co-regulated with *KDELR3* expression. Gene Set Enrichment Analysis (GSEA, FDR < 0.0001) of *KDELR3* co-expressed genes in TCGA skin cutaneous melanoma patients (cBioPortal)^30, 31^, revealed Gene Ontology (GO) term enrichment of ECM and trafficking pathways (consistent with previous data, Table 1, Supplementary Fig 2a), and pathways involved in the ERSR and response to unfolded proteins (Supplementary Fig. 6a). Quantitative mass spectrometry was used to analyze whole cell lysates of *KDELR3* knockdown compared to non-targeting controls and parental controls; GSEA analysis revealed the top-scoring, most consistent pathway using GO term enrichment showed upregulation of ER lumen proteins (Supplementary Fig. 6b). Enriched proteins included protein chaperones, lectins, and enzymes involved in protein folding and targeting misfolded proteins for degradation (including UGGT, ER Lectin, FKBP7, Calumenin), which is consistent with an increase in misfolded protein load in *KDELR3* knockdown cells^32^ We therefore asked how *KDELR3*’s role in the ERSR response is associated with its metastasis phenotype. Metastasis is known to be linked with ER stress, activating the UPR and therefore downstream signaling events that function to alleviate this stress^17^. High doses of ER stress, or an ineffective UPR have been associated with deleterious signals and ultimately cell death. We therefore hypothesized that one role of *KDELR3* in metastasis would be to alleviate ER stress-induced deleterious signaling (Supplementary Fig. 6c). We observed in four independent mouse models of melanoma (*N* = 6-13 mice per model) that *Perk* (*Eif2ak3*) transcription was negatively correlated with *Kdelr3* transcription (Fig. 4a), whereas *Gadd34* (*Ppp1r15a*) transcription was positively correlated (Fig. 4b). As PERK is a protein kinase and GADD34 a protein phosphatase that both act on EIF2α^33^, we hypothesized that KDELR3^−low^ cells are primed to activate the PERK-EIF2α arm of the UPR. We knocked down *KDELR3* (KD) in both 1205Lu and WM-46 human cell lines (shRNA knockdown; Supplementary Fig. 5b) and found that loss of *KDELR3* expression resulted in increased PERK and EIF2α protein levels in untreated cells, corroborating our mouse model data (Fig. 4c). We also saw a concomitant increase in PERK and EIF2α phosphorylation, suggesting constitutive activation of the PERK-EIF2α axis in untreated KD cells (Fig. 4c). The other two branches of the UPR pathways, the IRE1-XBP1 and ATF6α axes, were inactive in untreated *KDELR3* KD cells (Supplementary Fig. 6d-e). Tunicamycin, a chemical inhibitor of N-glycosylation that induces ER stress in cells, was used as a positive control (Fig. 4c; Supplementary Fig. 6d-e).

**Figure 4:**
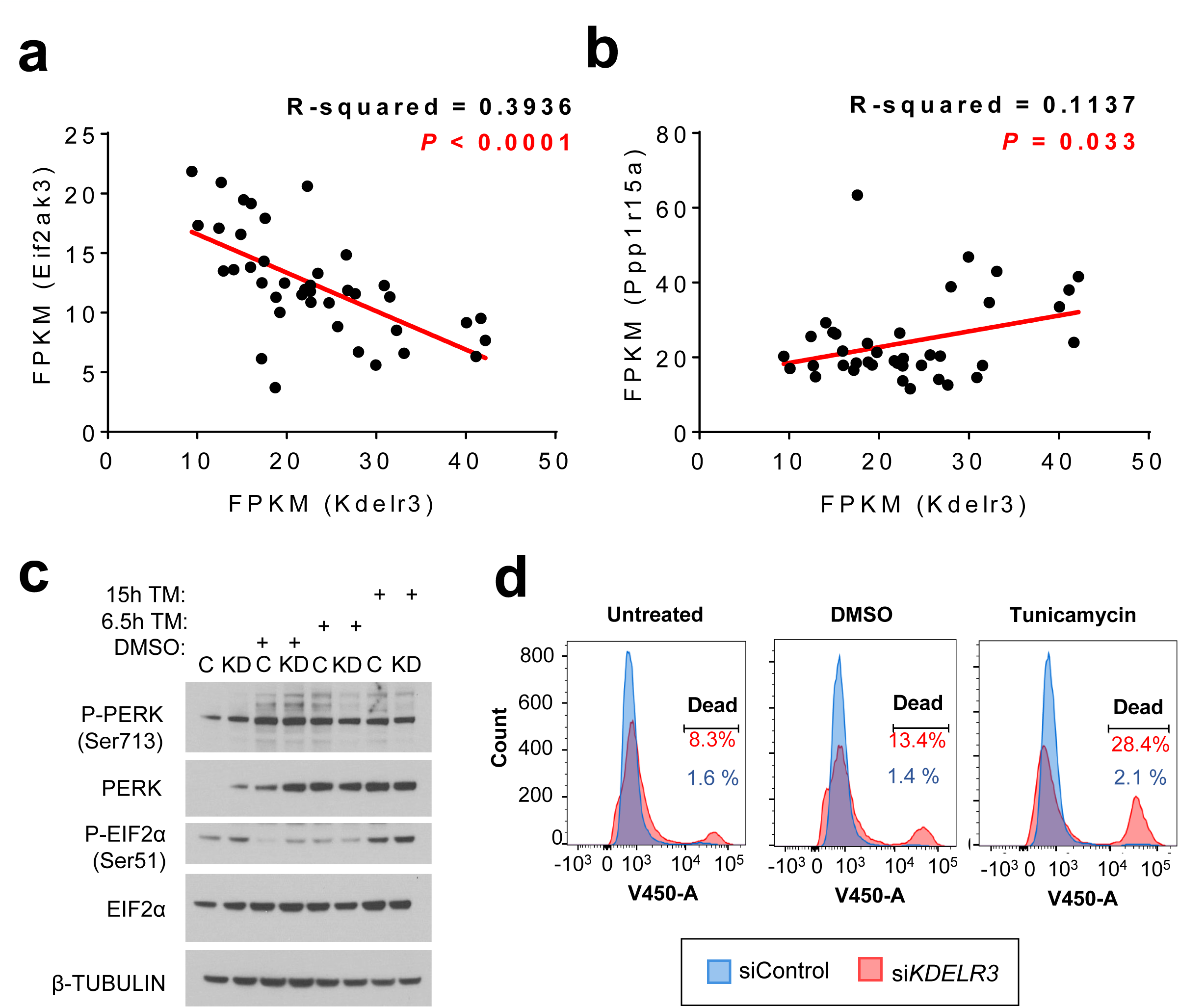
*KDELR3* and the ER Stress Response in metastatic melanoma. **a**, Scatter plot of *Eif2ak3* (*Perk*) RNA expression versus *Kdelr3* RNA expression. Linear regression analysis, R-squared = 0.3936, *P* < 0.0001. **b**, Scatter plot of *Ppp1r15a* (*Gadd34*) RNA expression versus *Kdelr3* RNA expression. Linear regression analysis, R-squared = 0.1137, *P* = 0.03. **a**, **b**, Data from four independent mouse models of melanoma (see methods). Each dot represents one mouse. M1, *N*= 9 mice; M2, *N*= 6 mice; M3, *N*= 12 mice; M4, *N*= 13 mice. **c**, Western blot analysis of PERK and eIF2α signaling (1205Lu cells). Non-targeting control (shControl; C) and *KDELR3* knockdown (sh*KDELR3*; KD) cells were untreated or treated with 3 µg/ml tunicamycin (TM) for the indicated time. Immunoblot with antibodies specified, β-Tubulin loading control. **d**, Live/dead violet cell stain in *KDELR3*-knockdown 1205Lu cells. Untreated, DMSO, and tunicamycin (2.5 μg/ml) treatment groups were treated 18 hours before collection. Right-hand peak on graph indicates percentage dead cells. Representative of 3 independent experiments.

Untreated *KDELR3* KD cells exhibited reduced levels of BiP, an essential protein chaperone necessary for activation of all arms of the UPR^17^, suggesting that retrograde transport in non-stressed cell may be required for long-term maintenance of BiP homeostasis (Supplementary Fig. 6e)^19^. These data indicate that loss of *KDELR3* expression disrupted ER homeostasis, resulting in a dysregulated UPR, which has previously been linked with ER stress-associated cell death^34^. We hypothesized that KDELR3 functions to alleviate deleterious ER stress-induced signaling (Supplementary Fig. 6c). To test this, we asked if *KDELR3* knockdown sensitizes metastatic melanoma cells to ER stress-induced death. We treated cells with tunicamycin, and measured cell death through flow cytometry using Live/Dead cell stain. We observed that siRNA-mediated knockdown of *KDELR3* expression resulted in a ∼5-fold increase in metastatic melanoma cell death over controls (8.3%, si*KDELR3*; 1.6%, siControl; Fig. 4d). These data suggest that *KDELR3* promotes cell survival in metastatic melanoma cells, which likely influences metastatic potential. *KDELR3*-knockdown cells have an enhanced sensitivity to ER stress induction with tunicamycin (>13-fold difference in cell death: 28.4%, si*KDELR3*; 2.1%, siControl; Fig. 4d). These data indicate that the ability of *KDELR3* to relieve ER stress is crucial for adaptation and survival in metastatic melanoma and may be instrumental to the metastatic phenotype.

### *KDELR3* mediates post-translational regulation of the metastasis suppressor KAI1

To further understand the role of *KDELR3* in metastasis, we queried if *KDELR3* knockdown would increase expression of known metastasis suppressors in melanoma. To address this, we screened protein expression of 5 melanoma metastasis suppressors following *KDELR3* knockdown^35, 36^. Of the 5 metastasis suppressors screened (BRMS1, Gelsolin, GAS1, NME1/NM23-H1, KAI1) only KAI1 demonstrated an increase in expression following *KDELR3* knockdown (Fig. 5a), in line with our hypothesis. Moreover, we observed a change in KAI1 molecular weight distribution following *KDELR3* knockdown, suggesting alterations in KAI1 post-translational modification. KAI1 protein upregulation was independent of transcriptional changes (Fig. 5b), supporting a regulatory role for KDELR3 at the post-translational level. KAI1 has been shown to influence metastasis through multiple mechanisms, including cell-cell adhesion, cell motility, cell death and senescence, and protein trafficking in many cancer types, including melanoma^37^. To further validate the role of KDELR3 on KAI1 protein regulation, we exogenously expressed KAI1 protein in 1205Lu metastatic melanoma cells (in which endogenous KAI1 expression is relatively low) and co-expressed both *KDELR3-001* and *KDELR3-002*. Corroborating our initial findings, we found that increased *KDELR3* expression resulted in dramatically reduced KAI1 protein levels (Fig. 5c), which could not be accounted for by KAI1 transcriptional changes (Fig. 5d-e). KAI1 protein glycosylation pattern was impacted reciprocally by knockdown and overexpression experiments, supporting the notion that KAI1 post-translational modification pathways are regulated by KDELR3, including an upregulation of a high molecular weight band in *KDELR3* knockdown cells (Fig. 5f, red arrow) that we showed corresponds to a highly glycosylated form of KAI1 (Fig. 5g). Glycosylated KAI1 has been linked to inhibition of cell motility and promotion of cell death^38^, and has been shown to influence N-cadherin clustering and bone metastasis in AML^39^.

**Figure 5:**
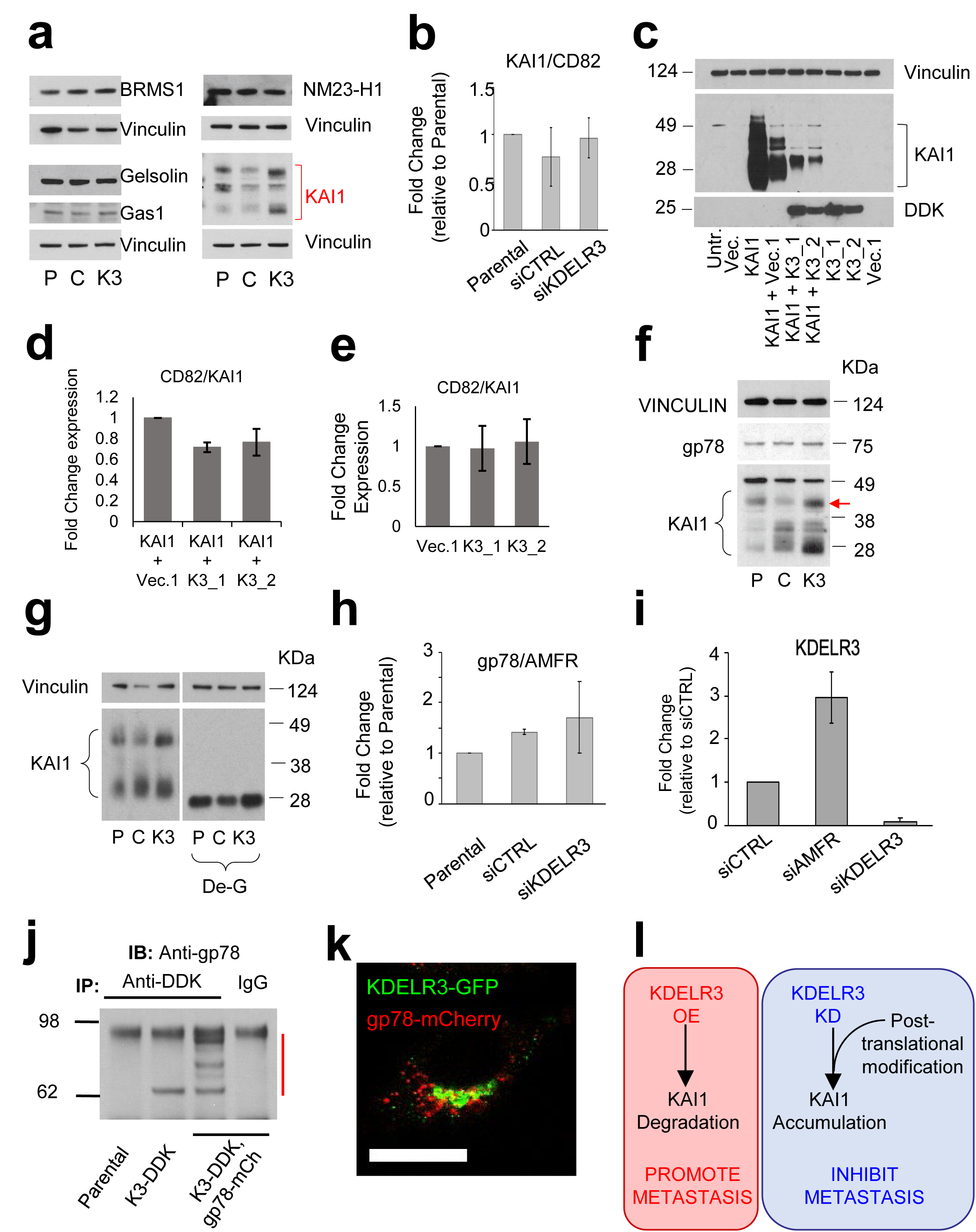
*KDELR3* regulates expression and processing of the metastasis suppressor KAI1. **a**, Screen of known melanoma metastasis suppressor expression following *KDELR3* knockdown (3 days post knockdown). P, Parental; C, siControl; K3, si*KDELR3.* **b**, qPCR of *KAI1* RNA expression (*CD82* gene) in siRNA knockdown cells (indicated), 3 days post knockdown. **c-e**, KAI1 protein (**c**) and RNA (**d**-**e**) expression in 1205Lu cells transfected with CD82/KAI1 overexpression (KAI1) or PCMV6-AC control vector (Vec.), *KDELR3* transcript 1with DDK tag (K3_1), *KDELR3* transcript 2 with DDK tag (K3_2), or PCMV6 control vector (Vec.1). Harvested 3 days post transfection. Equal protein amounts subjected to immunoblot analysis with an anti-KAI1 and anti-DDK antibody and anti-VINCULIN loading control (**c**). **f**, 1205Lu cells parental (P), and 1205Lu cells transiently transfected with control siRNA (C), and *KDELR3* siRNA (K3), harvested 3 days post transfection and equal protein amounts subjected to immunoblot analysis with an anti-KAI1 and anti-gp78 antibody. **g**, KAI1 protein expression in siRNA knockdown (indicated) 1205Lu cells harvested 3 days post transfection and treated with de-glycosylation enzymes (De-G). **h**, qPCR of gp78 RNA expression (*AMFR* gene) in siRNA knockdown cells (indicated), 3 days post knockdown. **f**, **g**, Anti-vinculin antibody used to control for protein loading. **i**, qPCR of *KDELR3* RNA expression in siRNA knockdown cells (indicated), 4 days post knockdown **j**, Co-immunoprecipitation of endogenous gp78 and mCherry tagged gp78 (gp78-mCh) with FLAG-tagged KDELR3 (K3-DDK) in stably transduced 1205Lu cells. Red line, gp78 expression. **k**, *pol2*>*KDELR3*-GFP (green) co-localizes with pol2>*gp78*-mCherry (red) in 1205Lu metastatic melanoma cells. Scale bars, 50 µm. **l**, Schematic of the KDELR3-KAI1 axis in melanoma metastasis. **a**-**f**, **h**, **j**-**k**, Representative of three independent experiments. **g**, **i**, Representative of two independent experiments.

Owing to our protein expression data, we hypothesized that KDELR3 regulates KAI1 protein degradation. We asked if KDELR3 regulates expression of the E3 ubiquitin ligase known to target KAI1, gp78/Autocrine Motility Factor Receptor^23, 40^, hereafter gp78. Although we saw no significant alterations in gp78 protein or RNA expression following *KDELR3* knockdown (Fig. 5f, h), we did observe a 3-fold increase in *KDELR3* transcription following gp78/*AMFR* knockdown, suggestive of a functional link between the two proteins (Fig. 5i). We identified a previously undescribed interaction between KDELR3 and gp78, which was supported by evidence of co-localization (Fig. 5j-k; Supplementary Fig. 7a). Interestingly, gp78 was first identified as a motility factor associated with metastasis in several cancers^41^, including melanoma. We asked if the KDELR3-gp78 interaction impacted gp78 function. We reasoned that gp78 ubiquitin ligase substrates would be upregulated following *gp78* knockdown, as these proteins would not be targeted for degradation; however, not all upregulated proteins identified will be direct gp78 substrates. Quantitative mass spectrometry was used to analyze whole cell lysates of gp78 (*AMFR*) knockdown or *KDELR3* knockdown cells compared to non-targeting controls. We could confirm that 43-57% of upregulated proteins matched between the gp78 and *KDELR3* knockdown groups. GSEA showed that the top-scoring, upregulated pathways (FDR <0.05) for both groups using GO term enrichment were those associated with the ER (Supplementary Table 3-4). This result suggests that both gp78 and KDELR3 act within similar cellular pathways and supports a role for KDELR3 in gp78 function, highlighting at least one mechanism through which KDELR3 can influence metastasis at the post-translational level. Since gp78 is a ubiquitin ligase known to function in ERAD, our data link KDELR3 to ERAD regulation. In summary, our work implicates KDELR3 in glycosylation of the metastasis suppressor, KAI1, and in its degradation through gp78 (and likely other ERAD effectors), thereby providing a mechanism for KDELR3’s influence on the metastatic phenotype (Fig. 5l).

### *KDELR3* correlates with late-stage metastasis and poor prognosis in melanoma patients

To assess how *KDELR3* contributes to melanoma progression in patients, we utilized multiple melanoma patient databases, The Cancer Genome Atlas^30, 42^ (TCGA) and Gene Expression Omnibus (GEO; GSE8401^29^, GSE19234^28^). We found increased expression of the *KDELR3-001* transcript, but interestingly not the alternate transcript, *KDELR3-002*, in late-stage (stage III and IV) metastatic melanoma patients compared to early-stage (stage I and II) melanoma patients (Fig. 6a), consistent with a role for *KDELR3* in melanoma progression. Metastatic melanoma patients with *KDELR3* copy number amplifications demonstrated reduced survival relative to patients without such alterations (Supplementary Fig. 7b). We next assessed melanoma patient survival using *KDELR3* expression as a prognostic marker (GEO^28, 29^). High *KDELR3*-expressing late-stage metastatic melanomas show statistically significant association with poor patient outcome, whereas *KDELR3* expression levels in early-stage primary tumor samples did not (Fig. 6b-c). Taken together these data strongly support a role for *KDELR3* in the advancement of late-stage metastatic melanoma and implicate *KDELR3* as a bona fide MetDev gene.

**Figure 6:**
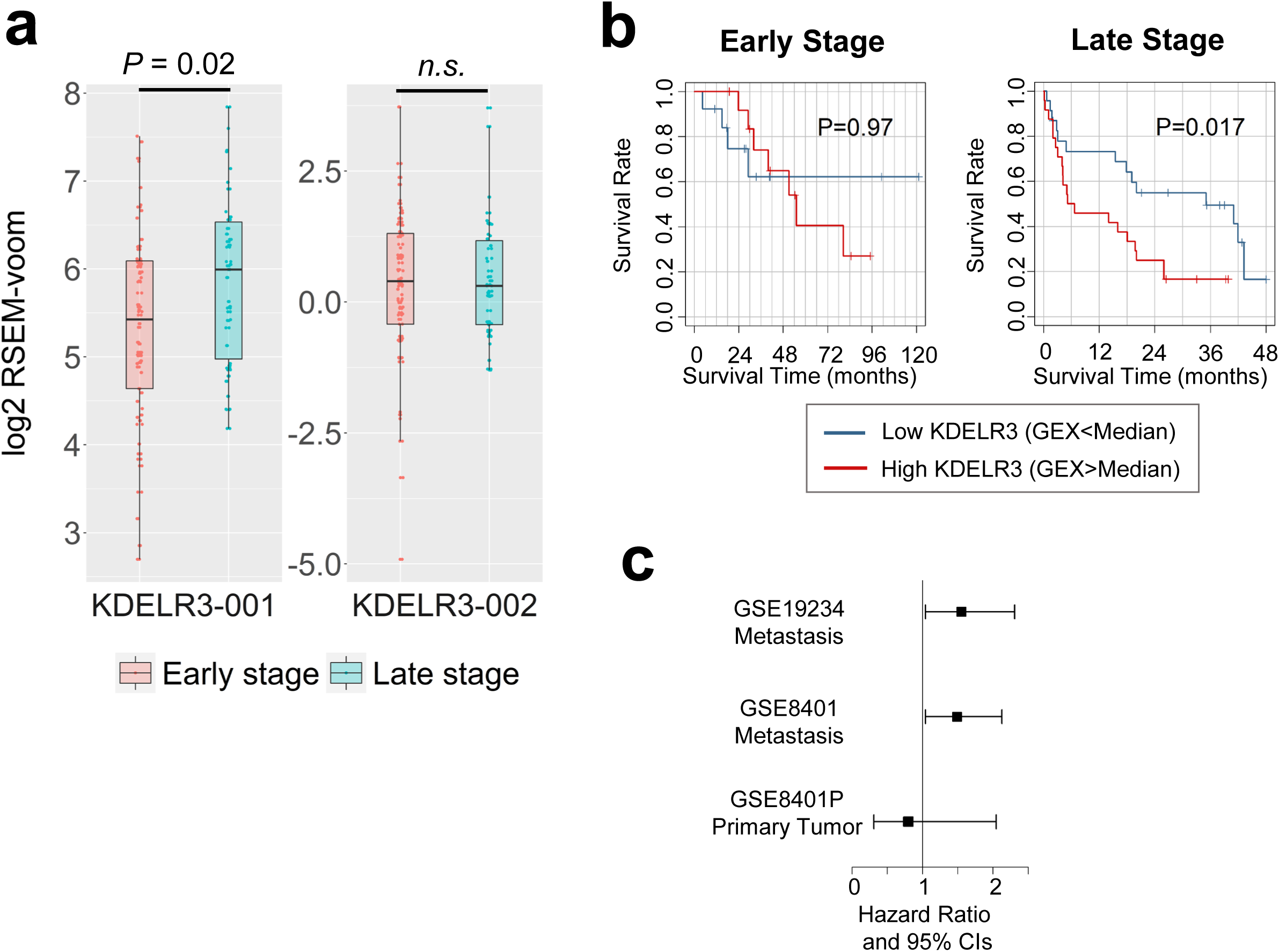
*KDELR3* expression correlates with advanced metastatic disease in patients. **a**, *KDELR3-001* and *KDELR3-002* patient expression data. Empirical Bayes moderated t-statistic (unpaired two-tailed test); *KDELR3-001*, ENST00000216014, *P* = 0.0202, *t* = 2.36, *df* = 102.17; *KDELR3-002*, ENST00000409006, *P* = 0.87, *t* = 0.16, *df* = 102.17. Boxplots of patient expression data from TCGA-SKCM dataset^30, 42^, depicting the 25th, 50th (median), 75th percentile, and extreme values of the transcript expression. “Early” stage (stages I/II, 62 patients). “Late” stage (stages III/IV, 39 patients). **b**, Kaplan-Meier estimated survival curves according to *KDELR3* expression in early-stage (GSE8401; n = 27, Stages I/II) and late-stage (GSE8401; n = 47, Stages III/IV) melanomas. Log-rank test. **c**, Association of *KDELR3* expression and survival in metastatic melanoma (GSE19234, n = 38; GSE8401, n = 47, Stages III/IV); HR = 1.62 (*P* = 0.028) and HR = 1.49 (*P* = 0.032) for GSE19234 and GSE8401, respectively. No significant association was found in the primary tumors (GSE8401, n = 27, Stages I/II); HR = 0.76 (*P* = 0.509). Cox regression model was used to test the association.

### *KDELR3* and *KDELR1* knockdown have opposing effects on lung colonization

As *KDELR3* is the only member of the KDELR family to be identified as a MetDev gene by our analyses, including embryonic-specific upregulation and consistent upregulation in melanoma cell lines, we posited that different KDELR members have different functions in melanoma etiology/progression. To address this, we asked which pathways were co-regulated with *KDELR1* expression and if these are the same or different relative to *KDELR3*-regulated pathways. GSEA analysis (FDR < 0.0001) of *KDELR1* co-expressed genes in TCGA skin cutaneous melanoma patients (cBioPortal)^30, 31^ revealed a strong enrichment of mitochondrial, metabolic and protein synthesis pathways (top 10 GO term enrichment, Fig. 7a), which differed from the most enriched pathways in *KDELR3* co-expressed genes that consisted predominantly of ECM, trafficking and ERSR pathways (top 10 GO term enrichment, Fig. 7b). Moreover, knockdown of *KDELR3*/*KDELR1* did not consistently alter expression of each other, suggesting that expression of these genes is not intrinsically linked (Supplementary Fig. 7c-d). These data intimate that *KDELR1* and *KDELR3* play different roles in melanoma progression. To test this, we compared the behavior of *KDELR3* and *KDELR1* knockdown cells using experimental metastasis assays. Notably, in contrast to *KDELR3* knockdown, which predictably diminished metastasis, *KDELR1* knockdown actually increased metastasis, suggesting that *KDELR1* contributes in a different way to melanoma etiology and can function as a metastasis suppressor (Fig. 7c-d). Moreover, analysis of *KDELR1* expression in skin cutaneous melanoma patients (TCGA) showed, unlike *KDELR3*, no significant difference between early-stage melanoma patients and late-stage metastatic melanoma patients (Fig. 7e). These data demonstrate that despite assumed redundancy between KDELR family members, *KDELR3* and *KDELR1* must have distinct roles, at least with respect to metastatic competence.

**Figure 7:**
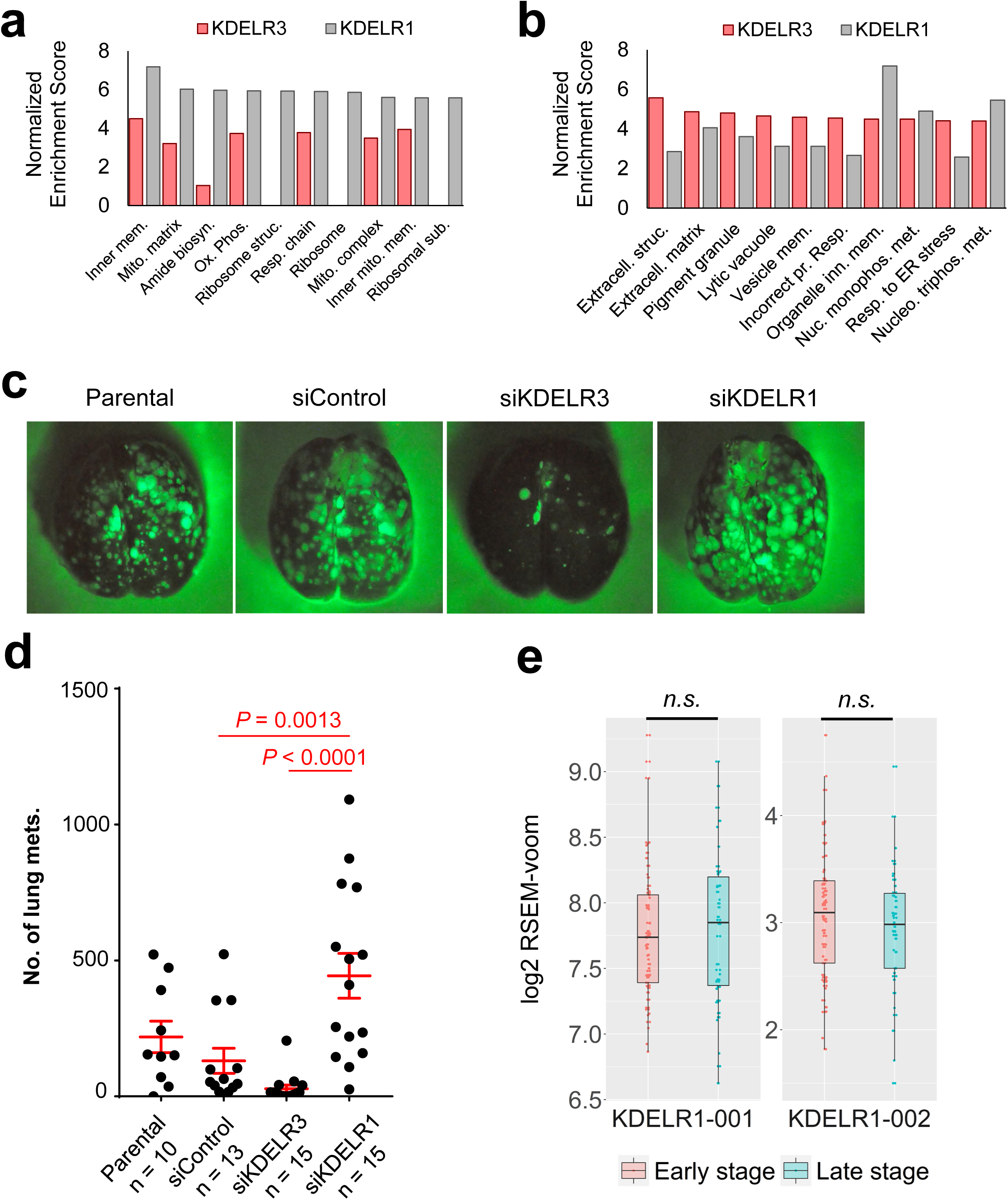
*KDELR1* knockdown increases lung colonization in tail vein metastasis assays. **a**, GSEA of gene co-expression within skin cutaneous melanoma patients of the TCGA (n = 479). Top 10 *KDELR1*-associated GO pathways represented, FDR < 0.0001. Go pathways in order: Organelle inner membrane; Mitochondrial matrix; Amide biosynthetic process; Oxidative phosphorylation; Structural constituent of ribosome; Respiratory chain; Ribosome; Mitochondrial protein complex; Inner mitochondrial membrane protein complex; Ribosomal subunit. **b**, GSEA of gene co-expression within skin cutaneous melanoma patients of the TCGA (n = 479). Top 10 *KDELR3*-associated GO pathways represented, FDR < 0.0001. Go pathways in order: Extracellular structure organization; Extracellular matrix; Pigment granule; Lytic vacuole; Vesicle membrane; Response to topologically incorrect protein; Organelle inner membrane; Nucleoside monophosphate metabolic process; Response to endoplasmic reticulum stress; Nucleoside triphosphate metabolic process. **c**-**d**, Tail vein metastasis of *KDELR1* siRNA-mediated knockdown human 1205Lu cells transduced with *Ferh-luc-GFP*. Parental, siControl and si*KDELR3* were used as controls. ANOVA with Tukey’s multiple comparisons test; siControl Vs si*KDELR1*, *P* = 0.0013, [mean difference (95% CI); −312.5(−521.7; −103.4)]; si*KDELR3* Vs si*KDELR1*, *P* <0.0001, [mean difference (95% CI); −415.3(−616.8; −213.7)]. *df* = 49, *F* = 10.8. CI, Confidence interval of differences. **e**, *KDELR1-001* and *KDELR1-002* patient expression data. Empirical Bayes moderated t-statistic (unpaired two-tailed test); *KDELR1-001*, ENST00000330720, *P* = 0.73, *t* = 0.35, *df* = 102.17; *KDELR1-002*, ENST00000597017, *P* = 0.39, *t* = −0.86, *df* = 102.17. Boxplots of patient expression data from TCGA-SKCM dataset^30, 42^, depicting the 25th, 50th (median), 75th percentile, and extreme values of the transcript expression. “Early” stage (stages I/II, 62 patients). “Late” stage (stages III/IV, 39 patients).

## Discussion

Here we propose that metastatic cancer cells exploit innate pathways that are hardwired within their cellular lineage to ensure proper development. These pathways, quieted in the differentiated cell, can be reactivated under pathologic conditions. The genetic/epigenetic reactivation of pathways that allow embryonic melanocytes to migrate, invade and colonize would represent an efficient strategy for melanoma cells to successfully metastasize. Here we employed a GEM model to identify, at the transcriptome level, novel genes that are required during melanocyte development and find that these are enriched in genes that are specific for progression of late-stage disease. We functionally validated 4 out of 4 genes tested, demonstrating the value of our dataset and supporting our hypothesis. We anticipate that other genes that passed our filtering criteria will ultimately prove to be functionally relevant and deserving of further analysis in future studies.

We report a mechanistic analysis of our top hit and melanoblast gene, *KDELR3*, a member of the KDEL receptor family. *KDELR3* has neither been previously associated with cutaneous melanoma metastasis nor investigated in depth in the literature. Differences between KDELRs have been cited in the literature but the main focus has been the role of KDELR1^19, 20, 43-45^. All three KDELR family members have been shown to mediate retrograde transport of proteins containing a C-terminus KDEL-like motif^19^. KDELRs typically reside in the cis-Golgi; however, tagged KDELRs are known to localize in both the cis- and trans-Golgi, which is consistent with our results^46^. Upon interaction with KDEL-like motif-containing proteins, KDELRs facilitate transport from the Golgi Apparatus back to the ER via COPI vesicles^47^. When this system fails, KDEL-like motif-containing proteins have been shown to be secreted out of the cell^19^. Our data demonstrating reduced BiP protein in stable *KDELR3* knockdown cells suggest that BiP is a genuine substrate for KDELR3 retrograde trafficking, and that without *KDELR3* expression melanoma cells are unable to maintain normal BiP levels. KDELRs appear to differ in the substrates that they preferentially transport, suggesting they have distinct roles within the cell^19^. How preferential substrate binding of KDELRs may affect cellular biology or disease etiology is still unknown.

Our study is the first to show that distinct KDELRs mediate dramatically different experimental metastasis phenotypes. We demonstrate that the embryonic melanoblast gene, *KDELR3*, is a metastasis enhancer in both mouse and human melanoma cells, whereas *KDELR1* suppresses metastasis, despite having extensive homology and similar retrograde trafficking functions. Our data allow a new perspective when interpreting existing KDELR literature and present a dichotomy between *KDELR3* and *KDELR1* metastasis phenotypes that could be leveraged in future studies to understand how these retrograde trafficking receptors function in disease. Moreover, Trychta and colleagues have reported tissue-specific KDELR expression patterns in rats implying that different KDELRs may have lineage-specific roles^19^. This is the first study to document *KDELR* expression in melanocyte development, and a specific role for *KDELR3*.

The KDEL receptors are intrinsically linked to ER stress and proteostasis. KDELR retrograde trafficking substrates include protein chaperones, protein folding chaperones, protein folding enzymes, enzymes that target proteins for degradation and glycosylation enzymes^19^. Cumulatively, these protein substrates help maintain correct protein processing, and regulate cellular response to ER stress^20^. However, the role of ER stress response in tumor progression has been much debated^48^. The success of proteasome inhibitors in the treatment of multiple myeloma patients^49^, as well as provocative data linking ER stress pathways to vemurafenib-resistant melanoma and immunotherapy sensitization, suggest UPR/ERAD biology could be harnessed for treating metastatic melanoma^50-54^. Our analysis implicates both UPR and degradation pathways of the ERSR as acting downstream of KDELR3. We show that *KDELR3* expression is critical for adaptation of melanoma cells to ER stress and provide evidence that PERK-EIF2α expression and activation is regulated by changes in *KDELR3* expression levels. Activation of the PERK-EIF2α pathway is known to result in translational attenuation, a cellular mechanism to alleviate ER load, causing translational rewiring of cells and affecting metastasis^15, 17, 48, 55-57^, which may contribute to KDELR3’s metastatic role.

We demonstrate that KDELR3 is a regulator of glycosylated KAI1, a tetraspanin glycoprotein with a well-documented metastasis suppressor role in tumors, including melanoma^23, 36, 37, 58, 59^. KAI1 functions at the cell membrane to mediate interactions between extracellular and intercellular signaling, which is key to its metastatic suppressor function. KAI1 glycosylation leads to changes in its membrane organization and therefore its ability to mediate this extracellular/intercellular signaling^38, 39, 60^. However, no studies have linked specific KAI1 glycosylated forms with its metastasis suppressor function *in vivo*. Our work notes specific glycosylated forms of KAI1 that are subject to KDELR3 regulation and associated with metastatic function. Future work would benefit from determining how critical each of these forms are to KAI1’s metastatic influence *in vivo*. Previously KDELR1 was shown to mediate signaling and transcriptional networks^43^, and at the protein level, in the relocation of lysosomes and modulation of autophagy^61^. However, KDELR3 was shown to be inactive in these processes. Here we link KDELR3 to post-translational regulation of protein, specifically through post-translational modification (glycosylation) and degradation of the metastasis suppressor, KAI1. Our data insinuate an interaction with gp78, implicating ERAD in this process. This biology may be informative for developing therapeutics for *KDELR3*-high metastatic melanoma patients.

We here identify an enrichment of ECM organization and trafficking genes within our MetDev cohort, consistent with a known role for these in metastasis^62-64^. Further analysis of these genes/pathways may prove a rich resource to uncover novel metastasis biology. We found that two such genes, *KDELR3* and *P4HA2* (a collagen prolyl 4-hydroxylase involved in ECM remodeling and associated with worse clinical outcome in melanoma patients^65^), from our 4-gene functional validation screen are tightly co-expressed in four independent mouse models and in human melanoma patients. This raises the possibility that expression of some genes within our MetDev cohort may be coordinated and/or networked to realize the complex and dynamic phenotypes exhibited by melanocytic cells during development and metastasis. Uncovering common upstream regulators of co-regulated genes could prove a powerful approach to target metastatic melanoma as multiple pathways could be targeted simultaneously.

To our knowledge this is the first example in which the mouse melanoblast transcriptome has been exploited to generate a resource of novel melanoma metastasis genes. The success of this study supports the use of developmental models to uncover innate melanoma biology that may be at the root of melanoma’s propensity to metastasize^2-9, 11-13, 66^. We anticipate that further exploration of *KDELR3* and other now-uncovered embryonic genes/pathways will facilitate the development of more effective treatment strategies for patients with advanced melanoma, and perhaps other tumor types. The field would further benefit from elucidation of the specific melanoblast cell characteristics/cell states that in fact contribute to metastasis. In summary, this work describes the creation of a novel resource of putative MetDev genes, enriched in genes that have functional roles in melanoma metastasis that may prove to be useful targets for designing more effective approaches to the treatment of melanoma patients.

## Methods

### Mouse models of melanoma

Experimental metastasis studies were performed using a filtered, single-cell suspension in PBS. 9.44×10^5^ (1205Lu) and 2×10^5^ cells (B16) were injected in 100 µl volume into the tail vein of 6-8-week-old ATHYMIC NCr-*nu*/*nu* mice (01B74, Frederick National Laboratory for Cancer Research) or C57BL/6N mice (Charles River, Frederick National Laboratory for Cancer Research), respectively. Lungs were removed from mice 4.5 weeks (1205Lu) or 24 days (C57BL/6N) post injection, perfused, and fixed in 10% phosphate buffered formalin (Fisher Scientific) for histology. Metastatic nodules were counted under a dissecting microscope.

Tumor growth studies were performed by injecting 3.47×10^5^ 1205Lu cells in a single-cell suspension subcutaneously into the flanks of 6-8-week-old ATHYMIC NCr-*nu*/*nu* mice (01B74, Frederick National Laboratory for Cancer Research). Tumor size was estimated using the formula: tumor volume (mm^3^) = 4/3π * (length/2) * (width/2) * height, where parameters were measured in mm.

Melanoblasts and melanocytes were isolated from the i*Dct*-GFP mouse model^8^. Embryonic development was timed based on number of days post-coitum. Pregnant females and newborn pups were placed on a doxycycline-enriched diet to activate expression of GFP.

Melanomas in Figure 4a-b and Supplementary Figure 3a were derived from the following four mouse melanoma models: **M1**; Albino C57BL/6 background, with *Braf*^CA/+^; *Pten*^flox/+^; *Cdkn2a*^flox/+^; *Tyr*-Cre^ERT2^-tg transgenic alleles. UV used as the tumor-inducing carcinogen; M1 mice were treated at postnatal day 3. **M2**; C57BL/6 background, with *Braf*^CA/+^; *Cdkn2a*^flox/+^; *Tyr*-Cre^ERT2^-tg; *Hgf*-tg transgenic alleles, UV used as the tumor-inducing carcinogen; M2 mice were treated at postnatal day 3. **M3**; C57BL/6 background, *Cdk4*^R24C^; *Hgf*-tg transgenic alleles, DMBA used as the tumor-inducing carcinogen; M3 mice were treated at postnatal day 3. **M4**; C57BL/6 background, with *Hgf*-tg transgenic allele, UV used as the tumor-inducing carcinogen; M4 mice were treated at postnatal day 3.

### Isolation of melanoblasts and melanocytes

FVB/N i*Dct*-GFP dams were fed doxycycline-fortified chow for the entire duration of gestation until collection of E15.5, E17.5 and P1 pups. Doxycycline was injected intraperitoneally at 80 μg/g body weight 24 hr before collection of P7 pups. A single cell suspension was generated from embryos and skin of newborn pups. Multiple litters were used for each developmental sage, and embryos/pups from each stage were pooled to ensure adequate numbers of GFP^+^ cells. The head was removed to prevent collection of GFP positive cells in the embryonic telencephalon, and melanocytes from the inner ear or from the retinal pigmented epithelium (RPE) were discarded. Excess tissue was also removed. The spinal cord was kept intact as some melanoblasts still remain in the neural crest area. At E17.5, P1 and P7 stages, most melanocytes have reached the dermis, thus only the skin was collected from these developmental stages. Back skin was immersed in a shallow layer of 1X PBS and subcutaneous fat was scraped off until skin appeared translucent. E15.5 was the youngest age assessed due to the necessity to capture sufficient cells for RNA-sequencing.

### Preparation of single cell suspensions

Tissue was minced and incubated for 30 min at 37°C in digestion media containing RPMI 1640 (Gibco Life Technologies) with 200 units/ml Liberase TL (Roche Applied Science). Up to 1g of tissue was digested per 5 ml digestion media. Tissue was processed using a Medimachine (BD Biosciences) and sterile medicon units (BD Biosciences). Cells were extracted using 1.5-2 ml of RFD solution (24 ml RPMI media, 6 ml FBS, 300 µl 5% DNase I) through a 20 ml syringe with 18-gauge needle. Collected cells were filtered through a 50-micron filter (BD Biosciences). This process was repeated until all the tissue was processed. Cells were spun at 1200 rpm, 4°C for 5 min and resuspended twice in a solution of 1% BSA in PBS and filtered through a 30-micron filter for sorting.

### Fluorescence activated cell sorting (FACS)

Embryos of the same developmental age that were heterozygous for the TRE-H2B-GFP gene but lacked the *Dct*-rtTA gene were used as negative controls. Cell doublets were excluded from the analysis. Cells were sorted based on GFP expression and SSC-A. Based on these reference sorts, gates were set so that background cells represented less than 10% of sorted cells.

### RNA Isolation and RNA sequencing

Cells were lysed in 10-fold TRIzol reagent (w/v), phases were separated by addition of 0.2X volume of chloroform, the aqueous phase was combined with an equal volume of 70% ethanol and applied to a RNeasy Micro column (Qiagen) and processed as per the manufacturer’s instructions. Paired-end sequencing libraries were prepared using 1 μg of purified RNA following the mRNA-Seq Sample Prep Kit according to the manufacturer’s instructions (Illumina). RNA-Seq libraries were sequenced on two lanes each of an Illumina GAIIx Genome Analyser to a minimum depth of 49 million reads. Sequence reads were aligned to the mm9 genome using the TopHat software (https://ccb.jhu.edu/software/tophat/index.shtml). Quantified Fragments Per Kilobase of transcript per Million mapped reads (FPKM) values were generated using the Cufflinks software (http://cole-trapnell-lab.github.io/cufflinks/). The UCSC KnownGenes gene models were used for guided alignment and quantification.

### Analysis of MetDev genes in patient survival

Based on the RNASeq data for the samples E15, E17, P1 and P7, we used DESeq2 to find differentially expressed genes comparing E15, E17 vs. P1, P7. We selected 467 up regulated genes with q-value < 0.1 (based on glm model) and log2FoldChange > 1.5. We then used the GEO dataset GSE19234 to perform survival analysis using Cox proportional hazards model for each gene. We to selected 43 genes that were correlated patient overall survival with the patient survival with p-value < 0.1 and HR >1. The Figure 1c-d showed the heatmaps of the gene expression (using z-score) for the 467 genes and 43 genes respectively. The sum of the total expression of the 43 genes forms the expression signature for prognosis prediction and the signature was tested on the new dataset GSE8401. Among the late-stage patients, the patients with high expression signature had significant poor survival compared to those with low expression (P=3.486e-5, Logrank test, Figure 1e) while for the early stage patients the two groups had no difference in survival (Figure 1e-f).

### Gene filtration pipeline for functional analysis

From our 467 identified melanoblast genes we first filtered for only those genes whose P7 expression level was minimal (FPKM < 2) i.e. no functional P7 gene expression to our knowledge, reasoning these would denote genes that truly had a unique role in melanoblast development compared to differentiated melanocytes. Next, we validated these by identifying the genes that are the intersect of the 467 genes with the differentially expressed genes from microarray expression data derived from our i*Dct*-GFP model (E17.5 vs P2 and P7)^21^. The microarray differential gene expression was identified using a linear regression model with contrast to compare embryonic versus postnatal stages and selected with a q-value < 0.1. The intersect yielded 233 genes. We acknowledge that the microarray data is not as thorough a representation of melanoblast/melanocyte development as our developmental cohort and therefore we may incur false negatives, we deemed this acceptable however to shorten our list for experimental validation. Next, we filtered the list to 81 genes with > 2.75 log2 fold increase expression in melanoblasts vs melanocytes and P-value <0.0003. Finally, we reasoned that genes with the greatest expression at embryonic stages would likely be the most functionally relevant, so selected for the top 20 greatest mean embryonic expression. Of these 20 we noted that 7 genes (*Kdelr3*, *P4ha2*, *Gulp1*, *Dab2*, *Lum*, *Aspn*, *Mfap5*) were all associated with Extracellular Matrix (ECM) or trafficking. Of these we chose to test the 3 least studied genes in metastasis (*Kdelr3*, *P4ha2*, *Gulp1*) to uncover novel metastasis biology, and the 1 gene well studied in metastasis (*Dab2*).

### Statistical analysis of KDELR3 expression in microarray data

Mouse developing melanoblasts (E17.5, n = 3) and differentiated melanocytes (P2, n = 3) were isolated and RNA extracted for microarray analysis as previously described^21^. The raw data (GSE25164 and unpublished, probe ID’s 1690129, 4920546) from Illumina mouseRef-8 v1.1 (GSM618249) expression beadchip were processed with variance stabilization transformation (VST) and quantile normalization as implemented in R *lumi* package (http://bioconductor.org/packages/release/bioc/html/lumi.html). Unpaired two-tailed t-test with Welch’s correction was used to compare the mean expression of *KDELR3* between the two developmental stages. As two probes for *KDELR3* on the Illumina beadchip showed high positive correlation (r = 0.987), the average *KDELR3* expression was analyzed.

### Analysis of The Cancer Genome Atlas (TCGA) skin cutaneous melanoma expression

All patient samples were collected between 0-14 days after disease classification (101 patients). Processed level 3 RNA-seq by Expectation-Maximization (RSEM) values^67^ were imported for melanoma patients from The Cancer Genome Atlas collection (TCGA-SKCM). Bioconductor *edgeR* (ttps://bioconductor.org/packages/release/bioc/html/edgeR.html) and *limma* (https://bioconductor.org/packages/release/bioc/html/limma.html) R packages were used for further processing and differential expression analysis. Transcripts with CPM (counts per million) greater than 1 in at least fifty percent of the samples were retained and processed with trimmed mean of M-values (TMM) and voom normalization methods^68^. The empirical Bayes moderated t-statistic test^69^ was applied to test the null hypothesis both for no difference in *KDELR3* expression, or for *KDELR1* expression level between early and late stage melanoma patients. A *P-*value of 0.05 or less was considered statistically significant.

### Statistics and general methods

All sample sizes were determined based on preliminary studies and prior knowledge of expected variability within assays. For animal studies, age-matched (6-8 weeks) female ATHYMIC NCr-*nu*/*nu* mice and C57BL/6 mice were randomly assigned to control and test groups. Blinding was used to quantify lung metastases counts. Where blinding was not used, data was analyzed using automated image analysis software when possible. All statistical tests used were deemed appropriate and met the assumptions required. Where necessary unequal variance was corrected for, or if no correction was used variation was assumed equal based on prior knowledge of the experimental assay. All cell lines used in this paper were identified correctly as per the International Cell Line Authentication Committee, version 8.0 (NB. MDA-MB-435 and MDA N cell lines in NCI60 were correctly identified as melanoma-derived cell lines). All cell lines used in experiments were screened for mycoplasma contamination and were tested negative for mycoplasma contamination. Cell lines were authenticated by examining their expression of melanoma markers using qPCR and RT-PCR analyses, and validating expression levels to those previously reported in published data. Human melanoma cell lines (1205Lu, WM-46 and SK-MEL-28) were validated using human-specific *TRP2*, *SOX10*, *TYRP-1* primers. Mouse melanoma cell line (B16) was validated using mouse-specific *MITF*, *TRP2*, *TYR* primers.

All mouse experiments were performed in accordance with Animal Study Protocols approved by the Animal Care and Use Committee (ACUC), NCI, National Institutes of Health. NCI is accredited by AAALACi and follows the Public Health Service Policy on the Care and Use of Laboratory Animals. Studies were carried out according to ASP # 16-007 and LMB-042. All animals used in this research project were cared for and used humanely according to the following policies: The U.S. Public Health Service Policy on Humane Care and Use of Animals (2015); the Guide for the Care and Use of Laboratory Animals (2011); and the U.S. Government Principles for Utilization and Care of Vertebrate Animals Used in Testing, Research, and Training (1985). The experimental records of animal studies in this project are maintained in a style consistent with ARRIVE guideline. Here we follow the guideline to report the results of animal studies in this manuscript.

### Code Availability

Upon acceptance of the manuscript custom code will be made publicly available and a full code availability statement will be included here.

### Melanoma cell lines

Human melanoma cells, 1205Lu and WM-46, were obtained from the Wistar Institute (courtesy of Meenhard Herlyn). SK-MEL-28 cells were obtained from (ATCC). 1205Lu cells were cultured in Tu2% media (as described by the Wistar Institute). WM-46 and SK-MEL-28 cells were cultured in 1X RPMI 1640, with 10% Serum and 2mM L-Glutamine (Gibco Life Technologies). For WM-46 cells flasks were coated with 0.1% gelatin (Stemcell). B16 mouse melanoma cells were obtained from Isaiah J. Fidler, M. D. Anderson Cancer Center^70^. Human 1205Lu cells were transduced with a high multiplicity of infection (MOI) of FerH-ffLuc-IRES-H2B-eGFP expressing lentivirus (11346-M04-653, Frederick National Laboratory for Cancer Research, Proteomics Facility, courtesy of Dominic Esposito)^70^. GFP-expressing cells were sorted using Fluorescence-activated cell sorting (BD FACSDiva 8.0.1, Flow Cytometry Core Facility, National Cancer Institute).

GIPZ^TM^ Lentiviral shRNA Particles were obtained from Dharmacon™. *KDELR3* shRNA (V3LHS_307898, gene target sequence: TGTGCCTATGTTACAGTGT), or non-silencing negative control (RHS4348) lentivirus were infected at both 34-43 transducing units (TU)/ cell, and also at 25 TU/cell for a separate experiment. Cells were selected and maintained in puromycin selection.

Wobble mutant cell lines were generated using the QuikChange II Site-Directed Mutagenesis Kit (Agilent Technologies). The *KDELR3* shRNA recognition sequence was edited (t210c_c213a_t216c_t219c_a222c) from Myc-DDK-tagged *KDELR3* transcript variant 1 construct (RC201571, OriGene). TOPO cloning was used to clone place this sequence into the Gateway cloning system and the pENTR L1/L2 plasmid was combined with C413-E19 pPol2 L4/R1 and pDEST-658 R4/R2 destination plasmids. Lentivirus was produced in the Protein Expression Laboratory, Leidos Biomedical Research, Inc., Frederick National Laboratory for Cancer Research. Cells previously transduced and selected with *KDELR3* shRNA and non-targeting control shRNA (Dharmacon, see previous), were transduced with 32.2 infection units (ifu) per cell, cells were transduced by spinoculation for 1 hour at 1200 xg. Infected cells were selected using blasticidin.

Forward primer

5’-GTAATGAAGGTGGTTTTTCTCCTCTGCGCATACGTCACCGTGTACATGATATATGGGAAATTCCG-3’

Reverse primer

5’-CGGAATTTCCCATATATCATGTACACGGTGACGTATGCGCAGAGGAGAAAAACCACCTTCATTAC-3’

Human gp78/*AMFR* expression vector was cloned using AMFR (NM_001144) sequence (RG209639, Origene) into pDest-653 destination vector by the Protein Expression Laboratory, Leidos Biomedical Research, Inc., Frederick National Laboratory for Cancer Research (mPol2p>Hs.AMFR-mCherry, 19771-M01-653). Lentivirus was produced in the Protein Expression Laboratory, Leidos Biomedical Research, Inc., Frederick National Laboratory. Cells were infected using a Multiplicity of Infection (MOI) of 5 and 8.8. Infected cells were selected using Fluorescence-Activated Cell Sorting for mCherry expression.

### siRNA knockdown of gene expression

For experimental metastasis assays siRNA knockdown experiments were performed 2 days prior to injection, as follows: siGENOME Human *KDELR3* (11015) siRNA SMARTpool (M-012316-02-0010, Dharmacon^TM^) for *KDELR3* siRNA knockdown in human cell lines, and siGENOME Mouse *Kdelr3* (105785) siRNA SMARTpool (M-052192-00-0005, Dharmacon^TM^) for *Kdelr3* knockdown in the mouse cell lines. For control knockdown, siGENOME Non-Targeting siRNA Pool #1 was used (D-001206-13-20, Dharmacon). For *KDELR1* knockdown in human cells, siGENOME Human *KDELR1* siRNA SMARTpool (M-019136-01-0005, Dharmacon^TM^) was used, Gene knockdown was done following the manufacturer’s instructions using DharmaFECT 1 Transfection Reagent (T-2001-02, Dharmacon). All other assays were performed using both the siGENOME siRNAs, including siGENOME Human *AMFR* siRNA SMARTpool (M-006522-01-0005, Dharmacon^TM^), and ON-TARGET Plus SMARTpool siRNAs for human *KDELR3* (L-012316-00-0005, Dharmacon^TM^), human *KDELR1* (L-019136-01-0005, Dharmacon^TM^) and ON-TARGET plus™ Control Pool (Non-targeting control, D-001810-10-20, Dharmacon^TM^). Using either the DharmaFECT 1 Transfection Reagent (T-2001-02, Dharmacon) or the Mirus TransIT-X2^®^ (Mirus) system as per the manufacturers’ instructions. Results were consistent between the all knockdown methodologies.

### Anchorage-independent growth assays

In 6-well plates 50,000 cells (B16 *Kdelr3* knockdown/ SK-MEL-28 *KDELR3* overexpression), 15,000 cells (WM-46 *KDELR3* knockdown), or 2,000 cells (WM-46 *KDELR3* rescue experiments) were plated in 0.4% Bacto™ Agar (Becton, Dickinson and Company) in 1X RPMI 1640 (Gibco Life Technologies) solution over a layer of 0.5% Agar-RPMI. Media was replenished twice weekly, and cell growth assessed at 4-weeks post plating. Wells were fixed in 10% Methanol/ 10% Acetic Acid fixation solution with subsequent staining using 0.01% crystal violet staining (Sigma-Aldrich)/ 10% Methanol solution. Colonies were analyzed under a dissecting microscope, and by imaging (Alpha Innotech imager) with subsequent analysis (Fluorchem HD2 software).

### Immunohistochemistry (IHC) and Immunofluorescence (IF) staining

Formalin-fixed paraffin-embedded (FFPE) immunofluorescence of i*Dct*-GFP mouse skin sections was performed using Heat Induced Epitope Retrieval (HIER) in Target retrieval buffer, pH 6 (Dako) for 7 min in an IHC microwave, followed by 15 min cooling on the bench. Overnight incubation (4⁰C) was with 1:50 Rabbit monoclonal KDELR3 (NBP1-00896, Novus Biological; 1DB_ID, 1DB-001-0000718990) followed by incubation with a biotinylated secondary antibody. Slides were blocked with rabbit serum prior to overnight incubation (4⁰C) with 1:500 PEP-8H (Dct) Rabbit monoclonal antibody^71^ (courtesy of Vincent Hearing). Avidin conjugated Dy-Light 594 (1:300) and Alexa Fluor secondary antibodies were used. Sections were analyzed using confocal microscopy.

FFPE lung sections were incubated for 15 min in Target retrieval buffer, pH6 (Dako) using HIER, and left for 15 min to cool. 1:50 KDELR3 (NBP1-00896, Novus Biological; 1DB_ID, 1DB-001-0000718990, Lot# CA36131)/ 1:400 KDEL Receptor 3 (L95) polyclonal (Bioworld Technology Cat# BS3124, RRID:AB_1663176, Lot# CA36131), and 1:250 HLA A (Abcam Cat# ab52922, RRID:AB_881225) antibodies were incubated for 1 hour, room temperature. Polymer detection was performed with ImmPRESS AP Reagent Kit, Anti-Rabbit Ig (Vector Laboratories). Chromagen staining was done using ImmPACT™ Vector Red Alkaline Phophatase substrate kit (Vector Laboratories), as per the manufacturer’s instructions. Sections were analyzed by a board-certified Veterinary Pathologist using the color deconvolution v9 algorithm in Aperio Image Scope v12.0.1.5027 software. Metastatic counts were generated with particle analysis in ImageJ software.

WM-46 cells stably transduced with FLAG-tagged *KDELR3-001* (ENST00000216014) were transfected with GFP-tagged AMFR construct (Origene, RG209639) using FuGENE^®^ HD transfection reagent as per the manufacturer’s instructions. Cells were plated on 0.1% gelatin-coated (Stemcell) glass coverslips and fixed with 4% paraformaldehyde for 15 min. Cells were permeabilized in 0.1% Triton-X-100 in PBS for 30 min, and blocked in 4% BSA in 0.05% Triton-X-100 in PBS for 10 min. Antibody incubation was for 1 hour, room temperature with: 1:50 Calnexin pAb (Abcam Cat# ab93355, RRID:AB_10563523, GR65788-22), 1:1000 Anti-DDK (FLAG) Clone 4C5 (OriGene Cat# TA50011-100, RRID:AB_2622345, Lot# A031), 1:75 GFP (D5.1) XP^®^ mAb (Cell Signaling Technology Cat# 2956S, RRID:AB_1196615, Lot# 2). Then co-stained with Alexa Fluor 488, 594 and 633 antibodies for 30 min, room temperature. Coverslips were mounted using mounting medium with DAPI (Vectashield, H-1200) and analyzed by confocal microscopy.

### Flow Cytometry Analysis of Melanoma Cells

Cell viability was assessed using LIVE/DEAD^TM^ Fixable Violet Dead Cell Stain kit (Invitrogen, Life technologies™). Three-days post siRNA knockdown of *KDELR3* or non-targeting control. Melanoma cells were fixed and stained as per the manufacturer’s instructions. When indicated, cells were treated with DMSO vehicle control or 2.5 μg/ml tunicamycin 18 hours before fixation Cell cycle analysis was performed using incubation of live cells with 10 µM 5-bromo-2’-deoxyuridine (BrdU) for 45 min (1205Lu) or 90 min (WM-46). Cells were fixed drop-wise with 100% ethanol to a final concentration of 70% ethanol at 4°C. Cells were resuspended in 0.5 mg/ml RNase A (37°C), and permeabilized with a solution of 5 M HCl 0.5% Triton-X-100 in dH2O for 20 min. Cells were incubated with 1:200 BrdU antibody (Cell Signaling Technology Cat# 5292S, RRID:AB_10548898, Lot# 3), and stained with either 1:200 Alexa Fluor-647 (1205Lu cells; Invitrogen), or 1:200 Alexa Fluor-488 (WM-46 cells; Invitrogen), then co-stained with 40 µg/ml PI solution.

### Reverse transcription and RT-PCR analysis of XBP1 splicing

Cultured cells were homogenized using TRIzol® reagent (Ambion^TM^) followed by vigorous agitation in chloroform, then spun at 12,000 × g, 15 min (4⁰C). The upper aqueous phase was utilized for RNA extraction using the RNeasy Mini Kit (Qiagen). Reverse transcription was carried out using the ImProm-II™ Reverse Transcription System (Promega) using Oligo (dT)20 oligonucleotides for poly-A tail detection. RT-PCR analysis of *XBP1* splicing was carried out using: *XBP1* F 5’-GGAGTTAAGACAGCGCTTGGGGA-3’ and *XBP1* R 5’-TGTTCTGGAGGGGTGACAACTGGG-3’ oligonucleotides and GoTaq® Green Master Mix (Promega), using a 58⁰C annealing temperature for 25 cycles. The reaction yields a 164 bp band (*XBP1*-unspliced) and a 138 bp band (*XBP1*-spliced). *GAPDH* loading control: *GAPDH*-F 5’-GGATGATGTTCTGGAGAGCC-3’, *GAPDH*-R 5’-CATCACCATCTTCCAGGAGC-3’.

### Real-Time quantitative PCR analysis of gene expression

SYBR Green dyes were used to run the reaction: GoTaq® qPCR Master Mix (Promega) with addition of CXR dye, or VeriQuest SYBR Green qPCR Master Mix (2X) (Affymetrix). Reactions were carried out according to the manufacturer’s guidelines on a 7900HT Fast Real-Time PCR system (Applied Biosystems) using SDS 2.4 software. 57°C/60°C annealing temperatures, 40 cycles were used. Oligonucleotides designed to detect cDNA of the 18S rRNA was used as a loading control for human cDNA: 18S-F 5’-CTTAGAGGGACAAGTGGCG-3’, 18S-R 5’-ACGCTGAGCCAGTCAGTGTA-3’. *Gapdh* loading control was used for qPCR of mouse cDNA: *Gapdh*-F 5’-CTGGAGAAACCTGCCAAGTA, *Gapdh*-R 5’-TGTTGCTGTAGCCGTATTCA-3’. Individual human genes tested: *KDELR3*-F 5’TCCCAGTCATTGGCCTTTCC-3’, *KDELR3*-R 5’-CCAGTTAGCCAGGTAGAGTGC-3’, *KDELR1*-F 5’-TCAAAGCTACTTACGATGGGAAC-3’, KDELR1-R 5’-ATTGACCAGGAACGCCAGAAT-3’, *KDELR2*-F 5’-GCACTGGTCTTCACAACTCGT-3’, *KDELR2*-R 5’-AGATCAGGTACACTGTGGCATA-3’, *KDELR3-001* F 5’-TGACCAAATTGCAGTCGTGT-3’, *KDELR3-001* R 5’-TCAGATTGGCATTGGAAGACT-3’. *AMFR*-F 5’-GGTTCTAGTAAATACCGCTTGCT-3’, *AMFR*-R 5’-TCTCACTCACTCGAAGAGGGC-3’.

### Exogenous expression studies

For exogenous over-expression of CD82 and KDELR3 genes the following expression plasmids were used: CD82 transcript variant 1 (NM_002231) Human Untagged Clone (Origene, CAT#: SC324395), pCMV6-AC Tagged Cloning mammalian vector with non-tagged expression (Origene, CAT#: PS100020), KDELR3 transcript variant 2 (NM_016657) Human Myc-DDK-tagged ORF Clone (Origene, CAT#: RC216726), KDELR3 transcript variant 1 (NM_006855) Human Myc-DDK-tagged ORF Clone (Origene, CAT#: RC201571), pCMV6-Entry Tagged Cloning mammalian vector with C-terminal Myc-DDK Tag (Origene, CAT#: PS100001). TransIT^®^-LT1 (Mirus Bio LLC.). Expression plasmids were transfected into 1205Lu human metastatic melanoma cells. Manufacturer’s guidelines were followed using a Reagent: DNA ratio of 3 µl TransIT^®^-LT1 Reagent per 1 µg DNA.

### Western blot analysis of protein expression

Cells were lysed with two methods: 1% Triton X-100 Buffer (50 mM Tris, pH 7.5, 150 mM NaCl, 1% Triton X-100, 10 mM iodoacetamide, phosphatase inhibitor cocktail 2 and 3 (Sigma-Aldrich) and cOmplete protease inhibitor (Roche), 50 µM MG132), or in RIPA lysis buffer (Sigma) with phosphatase inhibitor cocktails 2 and 3 (Sigma) and cOmplete™ Protease Inhibitor Cocktail (Roche) as per manufacturers’ guidelines. Protein lysates were denatured in LDS sample buffer (Invitrogen) and sample reducing agent containing DTT (Invitrogen) at 70°C for 10 min, then run on a 4-12% Bis-Tris NuPAGE gel (Novex by Life Technologies) in MES SDS running buffer (Invitrogen). Nitrocellulose membranes were probed with the following antibodies: anti-PERK Phospho (Ser713) Antibody (BioLegend Cat# 649402, RRID:AB_10640071, Lot# B203140), and Cell Signaling antibodies: anti-eIF2α (Cell Signaling Technology Cat# 9722, RRID:AB_2230924, Lot# 13), Phospho-eIF2α (Ser51) (D9G8) XP^TM^ Rabbit mAb (Cell Signaling Technology Cat# 3398, RRID:AB_2096481, Lot# 6), PERK (D11A8) Rabbit mAb (Cell Signaling Technology Cat# 5683S, RRID:AB_10831515, Lot#5), ATF-6 (D4Z8V) Rabbit mAb (Cell Signaling Technology Cat#65880, Lot# 1), BiP (C50B12) Rabbit mAb (Cell Signaling Technology Cat# 3177S, RRID:AB_2119845, Lot# 8), β-Tubulin (9F3) Rabbit mAb (Cell Signaling Technology Cat# 2128, RRID:AB_823664, Lot# 7). For immunoblotting, the rabbit monoclonal antibody to CD82 (D7G6H) was used (Cell Signaling Technology Cat#12439S). The rabbit antibody to AMFR was used (Cell Signaling Technology Cat#9590, RRIS: AB_10860080). Anti-VINCULIN mouse mAb (Sigma Aldrich, Cat# V9131). GAS1 Rabbit Polyclonal Ab (Origene Cat# AP51781PU-N, Lot# SH08D402D), NME1/NDKA (NM23-H1) Rabbit antibody (Cell Signaling Cat# 3345, Lot# 1), Gelsolin (D9W8Y) Rabbit mAb (Cell Signaling Cat# 12953, Lot#1), BRMS1 Rabbit polyclonal antibody (Invitrogen, Cat# PA5-78885, Lot# U82788252).

### Mass spectrometry

Cell lysates were extracted 4 days post siRNA knockdown of *KDELR3*, *AMFR* or non-targeting control (siGENOME) using Dharmafect #1 transfection reagent. Cell lysates (250 μg each) were digested with trypsin using the filter-aided sample preparation (FASP) protocol as previously described with minor modifications^72^. Lysates were first reduced by incubation with 10 mM DTT at 55 °C for 30 min. Each lysate was then diluted with 8 M urea in 100 mM Tris-HCl (pH 8.5) (UA) in a Microcon YM-10 filter unit and centrifuged at 14,000 × g for 30 min at 4°C. The lysis buffer was exchanged again by washing with 200 μL UA. The proteins were then alkylated with 50 mM iodoacetamide in UA, first incubated for 6 min at 25 °C and then excess reagent was removed by centrifugation at 14,000 × g for 30 min at 4°C. Proteins were then washed 3 × 100 μL 8 M urea in 100 mM Tris-HCl (pH 8.0) (UB). The remaining urea was diluted to 1 M with 100 mM Tris-HCl pH 8 and then the proteins were digested overnight at 37°C with trypsin at an enzyme to protein ratio of 1:100 w/w. Tryptic peptides were recovered from the filter by first centrifugation at 14,000 × g for 30 min at 4°C followed by washing of the filter with 50 μL 0.5 M NaCl. The peptides were acidified and desalted on a C18 SepPak cartridge (Waters) and dried by vacuum concentration (Labconco). Samples analyzing the effect of *KDELR3* siRNA treatment alone were dimethyl labeled, as described, with the label being rotated between replicates^73^. Samples analyzing the effect of *KDELR3* or *AMFR* siRNA knockdown were quantitated using label-free methods. Dried peptides were fractionated by high pH reversed-phase spin columns (Thermo). The peptides from each fraction were lyophilized, and dried peptides were solubilized in 4% acetonitrile and 0.5% formic acid in water for mass spectrometry analysis. Each fraction of each sample was separated on a 75 µm × 15 cm, 2 µm Acclaim PepMap reverse phase column (Thermo) using an UltiMate 3000 RSLCnano HPLC (Thermo) at a flow rate of 300 nL/min followed by online analysis by tandem mass spectrometry using a Thermo Orbitrap Fusion mass spectrometer. Peptides were eluted into the mass spectrometer using a linear gradient from 96% mobile phase A (0.1% formic acid in water) to 35% mobile phase B (0.1% formic acid in acetonitrile) over 240 minutes. Parent full-scan mass spectra were collected in the Orbitrap mass analyzer set to acquire data at 120,000 FWHM resolution; ions were then isolated in the quadrupole mass filter, fragmented within the HCD cell (HCD normalized energy 32%, stepped ± 3%), and the product ions analyzed in the ion trap.

The mass spectrometry data were analyzed and either dimethyl labeling or label-free quantitation performed using MaxQuant version 1.5.7.4^74, 75^ with the following parameters: variable modifications - methionine oxidation and N-acetylation of protein N-terminus; static modification – cysteine carbamidomethylation; first search was performed using 20 ppm error and the main search 10 ppm; maximum of two missed cleavages; protein and peptide FDR threshold of 0.01; min unique peptides 1; match between runs; label-free quantitation, with minimal ratio count 2. Proteins were identified using a Uniprot human database from November 2016 (20,072 entries). Statistical analysis was performed using Perseus version 1.5.6.0^76^. After removal of contaminant and reversed sequences, as well as proteins that were only quantified in one of the three replicate experiments, the quantitation values were base 2 logarithmized and non-assigned values were imputed from a normal distribution of the data. Statistically significant differences were assigned using a two-way t-test with a p-value cut-off of 0.05.

### Protein de-glycosylation

1205Lu cells were transfected with control or *KDELR3* siRNAs as previously described in Methods. After 4 days cells were lysed in in 1% Triton X-100 Buffer (50 mM Tris, pH 7.5, 150 mM NaCl, 1% Triton X-100, 10 mM iodoacetamide, phosphatase inhibitor cocktail 2 and 3 (Sigma-Aldrich) and cOmplete protease inhibitor (Roche), 50 µM MG132). Lysates were diluted 1:2 with dH2O to minimize lysis buffer effect. 10 µl Deglycosylation Mix Buffer 2 (New England Biolabs) was added to 17 µg of protein a 40 µl total volume, samples were heated at 75°C for 10 minutes. After cooling 5 µl Protein Deglycosylation Mix II (New England Biolabs) was mixed in gently. Reaction was incubated at 37°C for 30 minutes before being transferred to 37°C for 1 hour. Reactions were analyzed by NU-PAGE Nu-PAGE and immunoblotted with the rabbit monoclonal antibody to CD82 (D7G6H) was used (Cell Signaling Technology Cat# 12439S).

### Co-immunoprecipitation

Cells were lysed in 1% Triton X-100 Buffer (50 mM Tris, pH 7.5, 150 mM NaCl, 1% Triton X-100, 10 mM iodoacetamide, phosphatase inhibitor cocktail 2 and 3 (Sigma-Aldrich) and cOmplete™ Protease Inhibitor Cocktail (Roche), 50 µM MG132). Clarified lysates were pre-cleared by incubation with Dynabeads Protein A (Thermo-Fisher Scientific), at 4°C for 30 min. 2mg of total proteins lysate were immunoprecipitated with Dynabeads protein A-antibody complexes, using an anti-gp78 or anti-DDK antibody and their respective IgG isotypes: rabbit IgG (BD Pharmingen) and mouse IgG (Santa Cruz). Incubation with rotation overnight at 4°C was performed. Immunoprecipitates were washed five times with washing buffer (50 mM Tris, pH 7.5, 150 mM NaCl, 0.1% Triton X-100) and were resuspended in 50 µl of elution buffer containing washing buffer, NuPAGE LDS sample buffer and NuPAGE sample reducing agent, mixed as per manufacturer’s instructions (Invitrogen). Proteins were analyzed by Nu-PAGE and immunoblotted using an enhanced chemiluminescence (ECL) method. For immunoblotting anti-DDK antibody (Origene TA50011) or (Ab2) to amino acids 579–611 of gp78 was used; this antibody was previously described^23^.

## Acknowledgements

This research was supported in part by the NCI Intramural Research Program of the NIH. PJM was also supported in part by the NCI Director’s Innovation Award. MRZ was supported in part by the following grant: NIH/NCI K22CA163799. TG was supported in part by the HHMI Research Scholars Program, Howard Hughes Medical Institute. HTM funded in part by the NIH Comparative Biomedical Scientist Training Program in partnership with University of Maryland, College Park, and the National Cancer Institute. We would like to thank the Dr. Meenhard Herlyn for providing melanoma cell lines included in this study. We are grateful to Cari Graff-Cherry for maintenance and care of mouse lines, Dr. Stephen Locket for microscopy analysis, Dr. Joe Kalen and Nimit Patel for imaging of developing mice, and Drs. Dominic Esposito and Carissa Grose for lentiviral vector preparation. We acknowledge Dr. Yves Pommier and William Reinhold for NCI60 data and CellMiner analyses. We thank Jennifer Dwyer, Shelley Hoover and Bih-Rong Wei from the Molecular Pathology Unit for slide scanning and IVIS imaging. We acknowledge Leidos at the British Columbia Cancer Agency for gifting us free RNA-sequencing services. We are also grateful to Dr. Lalage Wakefield for useful discussions.

## Author Contributions

KLM, GM, wrote the manuscript. GM, KLM, PJM, AS, AMM, CPD, AMW, YCT, HTM, SD, PSM, LMJ participated in experimental/ study design. KLM, PJM, AS, AMM, HTM, TG, YCT, MRZ, EPG, SD, LMJ generated the experimental data. GM, KLM, PJM, AS, AMM, MPL, HHY, HTM, YCT, AMW, EPG, CPD, HA, SD, PSM contributed intellectually to the work. MPL, HHY, AMM, SD performed bioinformatic and statistical analysis of data.

## Data Availability

Upon acceptance of the manuscript all data will be made publicly available and a full code availability statement will be included here.

