## Supplementary figures and tables for "Melanoblast transcriptome analysis reveals novel pathways promoting melanoma metastasis"

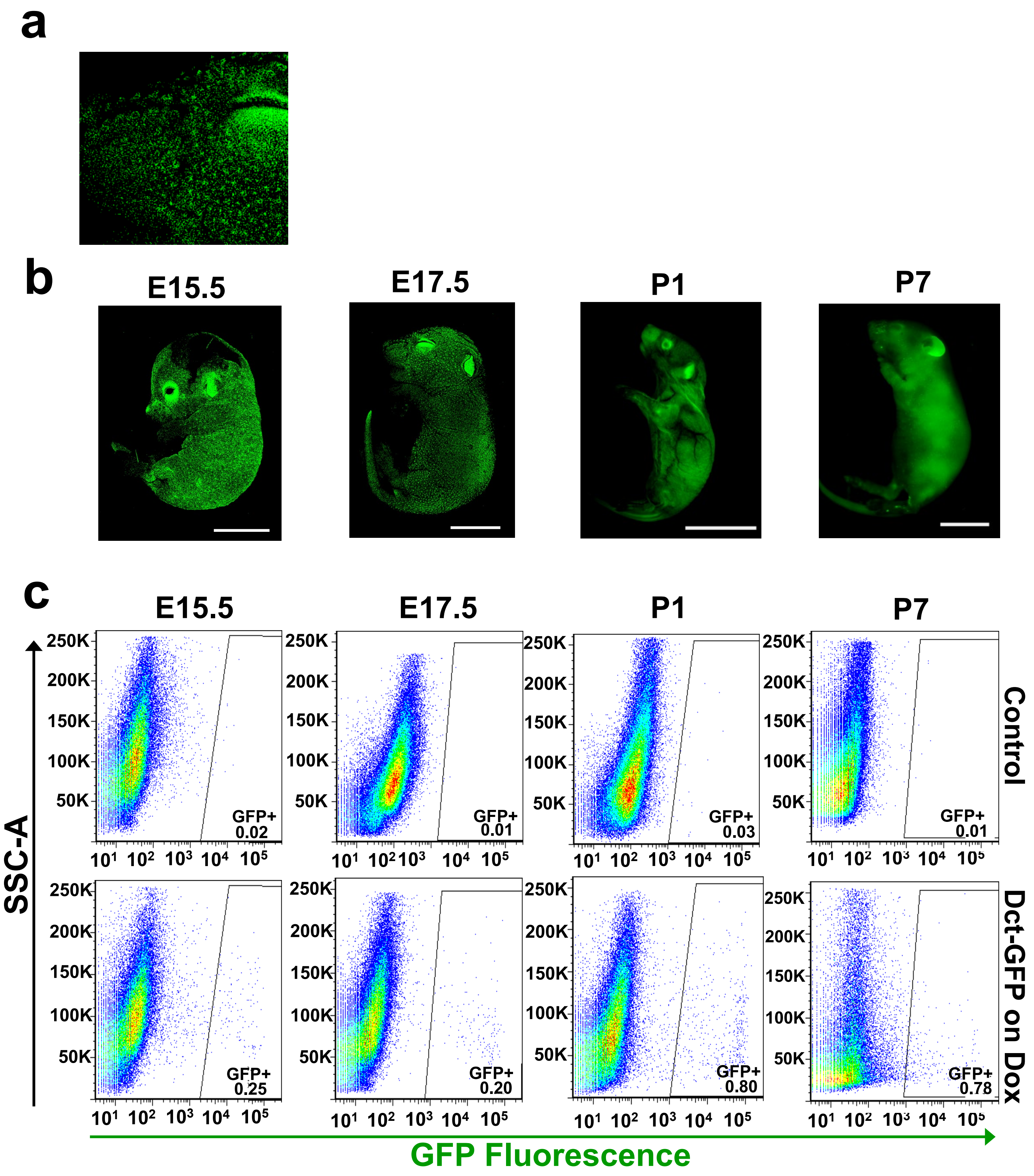
a
b
E15.5
E17.5
P1
P7
c
E15.5
E17.5
P1
P7
SSC-A
Control
Dct-GFP on Dox
GFP+ 0.02
GFP+ 0.01
GFP+ 0.03
GFP+ 0.01
GFP+ 0.25
GFP+ 0.20
GFP+ 0.80
GFP+ 0.78
250K
200K
150K
100K
50K
$10^1$ $10^2$ $10^3$ $10^4$ $10^5$
GFP Fluorescence

**Supplementary Figure 1: Characterizing the mouse melanoblast transcriptome. a-b**, Confocal imaging of i*Dct*-GFP embryos and pups. Zoom in of Embryonic day 17.5 pup showing melanoblasts localized to hair follicles on face (**a**). Embryonic day 15.5 and 17.5 (E15.5 and E17.5 respectively; **b**). Postnatal day 1 and postnatal day 7 (P1, P7 respectively; **b**). Magnification E15.5 and E17.5 is x5, scale bars, 5 mm; magnification P1 and P7 is x1. Scale bars, 10 mm. **c**, FACS-sorting of GFP-positive cells from *i*Dct-GFP mice. SSC-A: Side scatter area. Proportion of GFP-positive cells indicated ($GFP^+$).

**a**

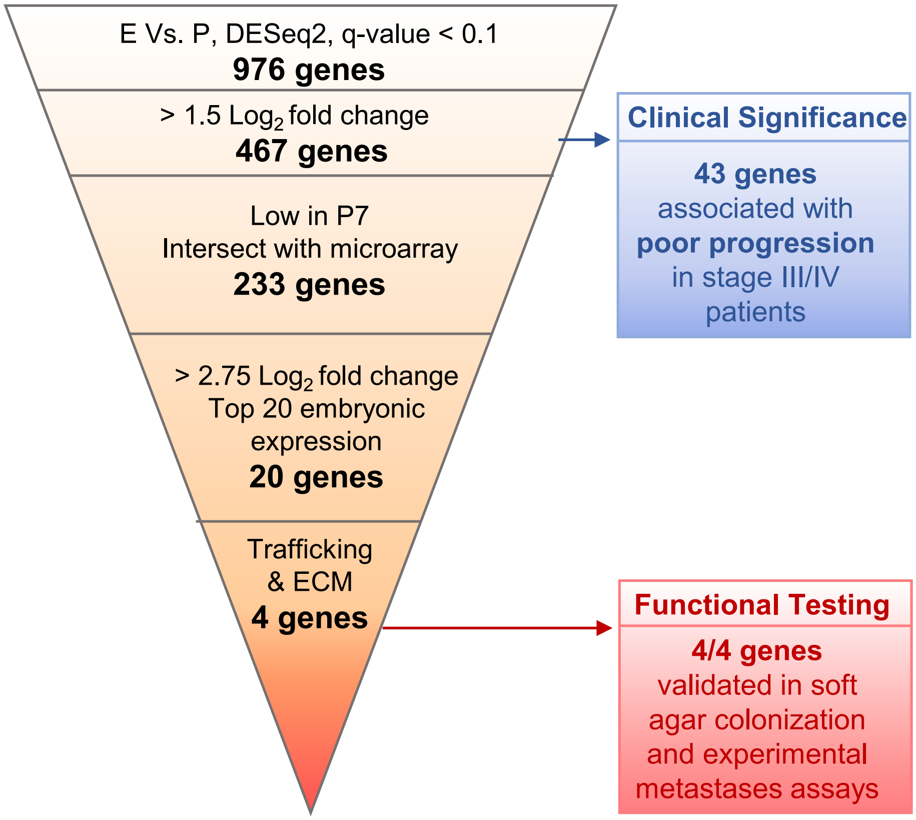
E Vs. P, DESeq2, q-value < 0.1
976 genes
> 1.5 Log2 fold change
467 genes
Low in P7
Intersect with microarray
233 genes
> 2.75 Log2 fold change
Top 20 embryonic expression
20 genes
Trafficking & ECM
4 genes
Clinical Significance
43 genes
associated with
poor progression
in stage III/IV
patients
Functional Testing
4/4 genes
validated in soft
agar colonization
and experimental
metastases assays

**Supplementary Figure 2: Validation of melanoblast genes in metastasis: workflow for gene prioritization. a**, Gene expression from Embryonic stages day 15.5 and 17.5 (E) was contrasted with gene expression from Postnatal stages day 1 and 7 (P) using DESeq2, with a False Discovery Rate (FDR) threshold of 0.1, q-value < 0.1 to yield 976 genes, of which 467 genes had >1.5 $Log_2$-fold change. Testing clinical significance of MetDev genes (blue box): Cox proportional hazards modeling of these 467 genes in training dataset GSE19234, yielded a 43-gene signature that was associated with poor progression in late-stage (stage III/IV) metastatic melanoma of both GSE19234 and GSE8401 (testing dataset). The 43-gene signature did not predict poor patient prognosis from early stage (stage I/II) primary tumors (GSE8401). Further prioritization of the 467 melanoblast genes took place purely on a statistical basis: All genes with a FPKM > 2 in P7 were filtered out. The intersect of embryonic genes to those determined in a separate microarray study of melanocyte development (E17.5 versus P2 and P7) using a linear regression model, selecting for q-value < 0.1, was determined. 233 genes were prioritized from the 467 gene list. The top 20 genes were determined by > 2.75 $Log_2$-fold change, P-value < 0.0003, and top 20 highest mean expression from embryonic stages. 4 genes were selected based on their role in trafficking and Extracellular Matrix (ECM), and their novelty in melanoma metastasis.

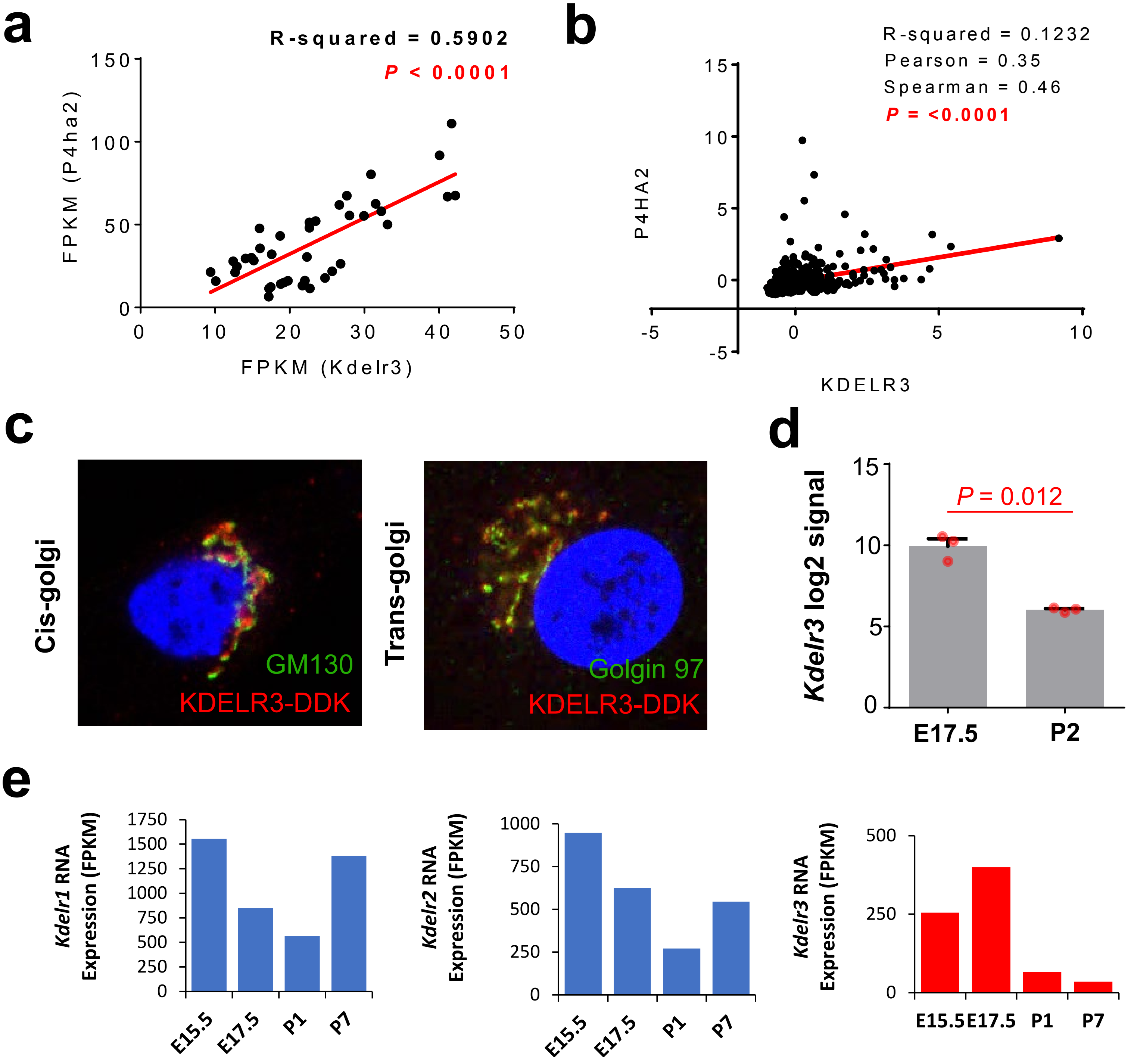

a
R-squared = 0.5902
P < 0.0001
FPKM (P4ha2)
FPKM (Kdelr3)
b
R-squared = 0.1232
Pearson = 0.35
Spearman = 0.46
P = <0.0001
P4HA2
KDELR3
c
Cis-golgi
GM130
KDELR3-DDK
Trans-golgi
Golgin 97
KDELR3-DDK
d
Kdelr3 log2 signal
P = 0.012
E17.5
P2
e
Kdelr1 RNA Expression (FPKM)
Kdelr2 RNA Expression (FPKM)
Kdelr3 RNA Expression (FPKM)
E15.5
E17.5
P1
P7

**Supplementary Figure 3: KDELR3 is an ER protein upregulated in melanoma. a**, Scatter plot of *P4ha2* RNA expression versus *Kdelr3* RNA expression in four independent mouse models of melanoma. Each dot represents one mouse. M1, *N*= 9 mice; M2, *N*= 6 mice; M3, *N*= 12 mice; M4, *N*= 13 mice. Linear regression analysis, R-squared = 0.5902, $P < 0.0001$. **b**, Scatter plot of *P4ha2* RNA expression versus *Kdelr3* RNA expression in human melanoma patients (cBioPortal, TCGA). Linear regression analysis, R-squared = 0.1232, $P < 0.0001$. Spearman = 0.46. Pearson = 0.35. **c**, Immunofluoresence in the 1205Lu human metastatic melanoma cell line. Flag-tagged KDELR3 expression (red) co-localizes with the cis-golgi marker, GM130 (green; left-hand panel) and trans-golgi maker, Golgin 97 (green, right-hand panel). Representative image of over 50 cells analyzed in three independent experiments. **d**, Independent study showing *Kdelr3* microarray expression data in mouse developing melanocytes[8], $P = 0.012$ n = 3, $t = 8.256$, $df = 2.12$. Unpaired two-tailed t-test with Welch's correction. **e**, RNA-seq data showing *Kdelr* family expression in mouse developing melanocytes**.**

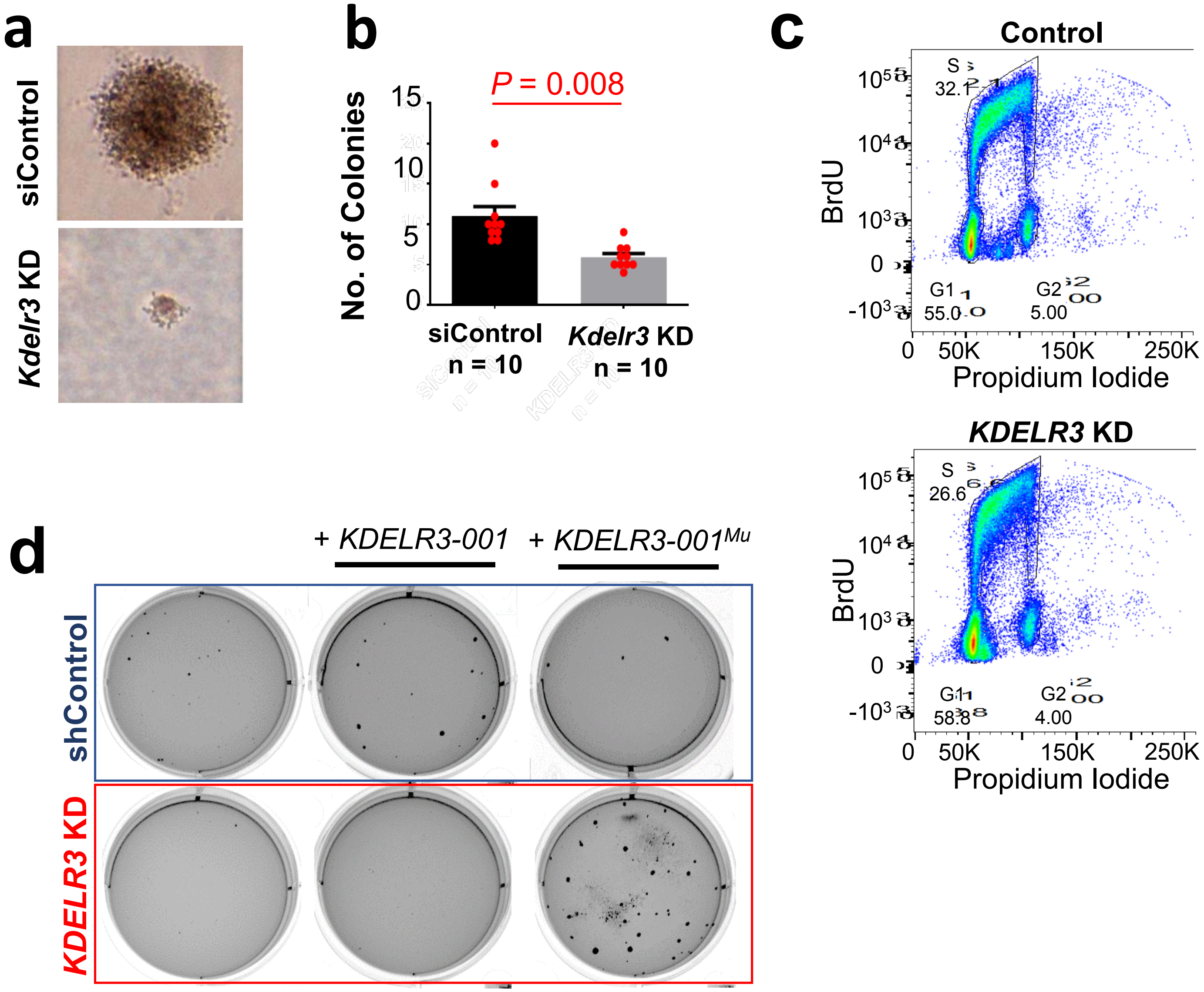


**Supplementary Figure 4: *KDELR3* mediates anchorage-independent growth in melanoma. a-b**, Soft agar colony formation assay with *Kdelr3* siRNA knockdown in mouse B16 cells versus non-targeting control, unpaired two-tailed student's t-test, $P$ = 0.008, $df$ = 18, $t$ =4.019. 10 wells analyzed per group. **c**, Flow cytometry analysis of BRDU incorporation (Alexa Fluor 488-A) and PI staining (DsRed-A) in WM-46 human melanoma cells. shRNA *KDELR3* knockdown does not affect the cell cycle, when compared to non-targeting control cells. G1-arrest, S-phase and G2-arrest populations were: 55 %, 32.1 %, and 5% in control cells, versus, 58.8 %, 26.8% and 4% in *KDELR3* knockdown cells. **d**, Rescue of soft agar colony formation in *KDELR3-001*$^{Mu}$ cells (WM-46).

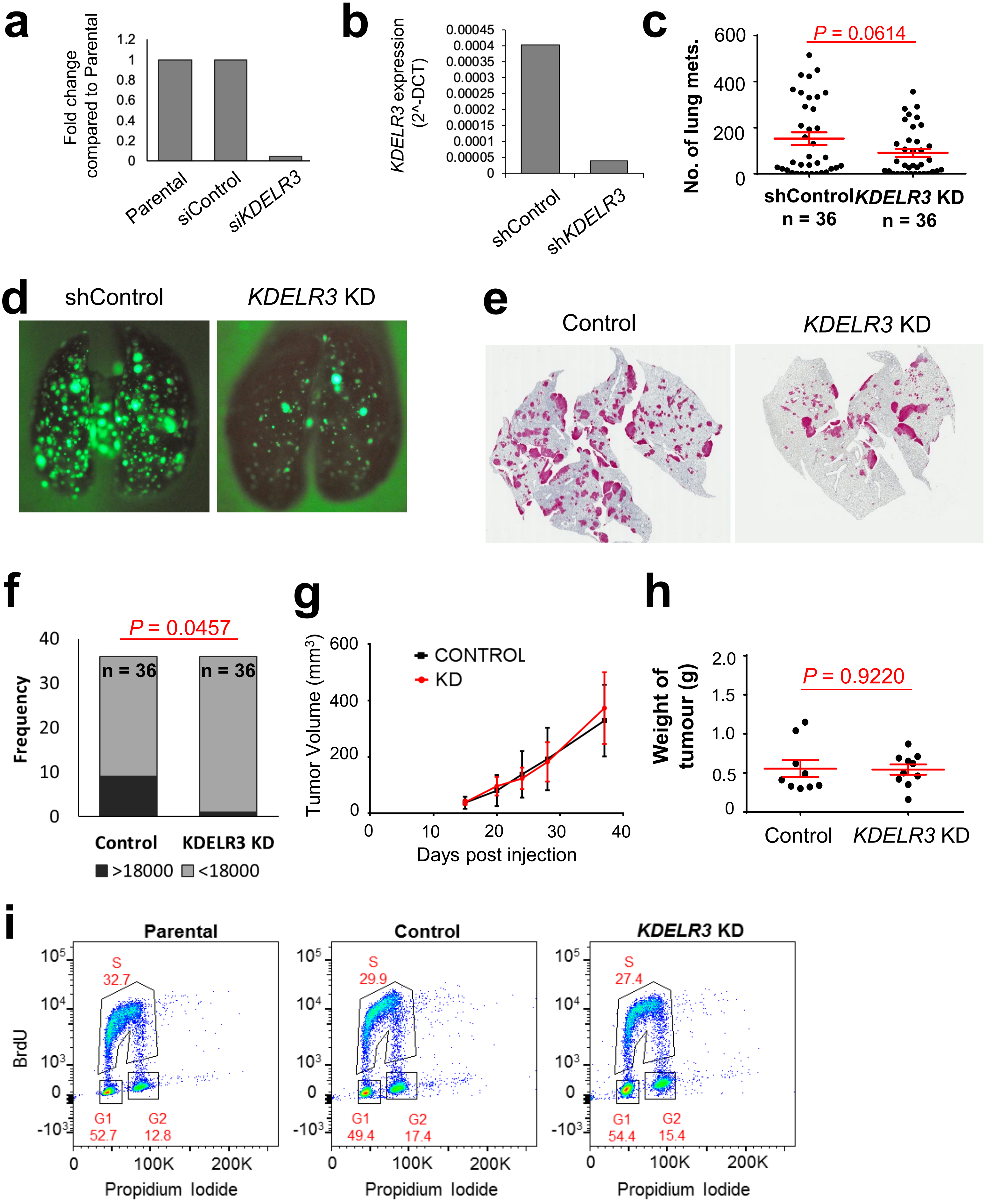

a
Fold change compared to Parental
1.2
1
0.8
0.6
0.4
0.2
0
Parental
siControl
siKDELR3
b
KDELR3 expression (2^-DCT)
0.00045
0.0004
0.00035
0.0003
0.00025
0.0002
0.00015
0.0001
0.00005
0
shControl
shKDELR3
c
No. of lung mets.
P = 0.0614
600
400
200
0
shControl n = 36
KDELR3 KD n = 36
d
shControl
KDELR3 KD
e
Control
KDELR3 KD
f
Frequency
P = 0.0457
40
30
20
10
0
n = 36
n = 36
Control
KDELR3 KD
>18000
<18000
g
Tumor Volume (mm³)
600
400
200
0
CONTROL
KD
0
10
20
30
40
Days post injection
h
Weight of tumour (g)
P = 0.9220
2.0
1.5
1.0
0.5
0
Control
KDELR3 KD
i
Parental
Control
KDELR3 KD
BrdU
10⁵
10⁴
10³
0
-10³
S 32.7
G1 52.7
G2 12.8
S 29.9
G1 49.4
G2 17.4
S 27.4
G1 54.4
G2 15.4
0
100K
200K
Propidium Iodide

**Supplementary Figure 5: *KDELR3* expression in melanoma cells enhances metastatic potential without affecting proliferation.** **a**, qPCR data of *KDELR3* expression in 1205Lu cells 3 days following siRNA knockdown of *KDELR3* (si*KDELR3*) or non-targeting control (siControl), as compared to Parental. **b**, qPCR data of *KDELR3* expression following shRNA knockdown of *KDELR3* (sh*KDELR3*) versus non-targeting control (shControl). **c-e**, Tail vein metastasis of *KDELR3* shRNA-mediated knockdown human 1205Lu cells transduced with *Ferh-luc-GFP*, (**c**) quantification of numbers of lung metastases. Unpaired two-tailed t-test with Welch's correction, $P = 0.0614$, $df = 59.21$, $t = 1.907$. **d**, Tail vein injection of *Ferh-luc-GFP*-transduced human 1205Lu cells[29] enables visualization of metastases (4 weeks post-injection). **e**, Human-specific HLA-A staining in FFPE sections (n = 36 per group); representative sections shown. **f**, Quantification of total area of metastases per section, determined by arbitrary units (ImageJ), two-tailed Fisher's Exact test, n = 36 lungs. **g**, Effect of *KDELR3* shRNA knockdown on subcutaneous tumor growth curve in 1205Lu cells, center and error bars represent mean ± standard deviation. Representative results from two independent experiments. **h**, Effect of *KDELR3* shRNA knockdown on subcutaneous tumor growth (1205Lu) represented by end tumor weight (40-days post injection). Representative of two independent experiments. **i**, Flow cytometry analysis of BrdU (Alexa Fluor-647) incorporation and Propidium Iodide staining in 1205Lu metastatic melanoma cells. shRNA *KDELR3* knockdown (*KDELR3* KD) does not affect the cell cycle, when compared to non-targeting control (Control) cells or parental cells. G1-arrest, S-phase and G2-arrest populations were: 52.7%, 32.7%, and 12.8% in parental cells; 49.4%, 29.9%, and 17.4% in non-targeting control (Control) cells; 54.4%, 27.4% and 15.4% in *KDELR3* knockdown cells. Representative of two independent experiments carried out in two different cell lines. **c-h**, Non-targeting shRNA control used. **d-f,** Cumulative data taken from three independent experiments. **c**, **h**, Lines and error bars depict mean ± s.e.m.

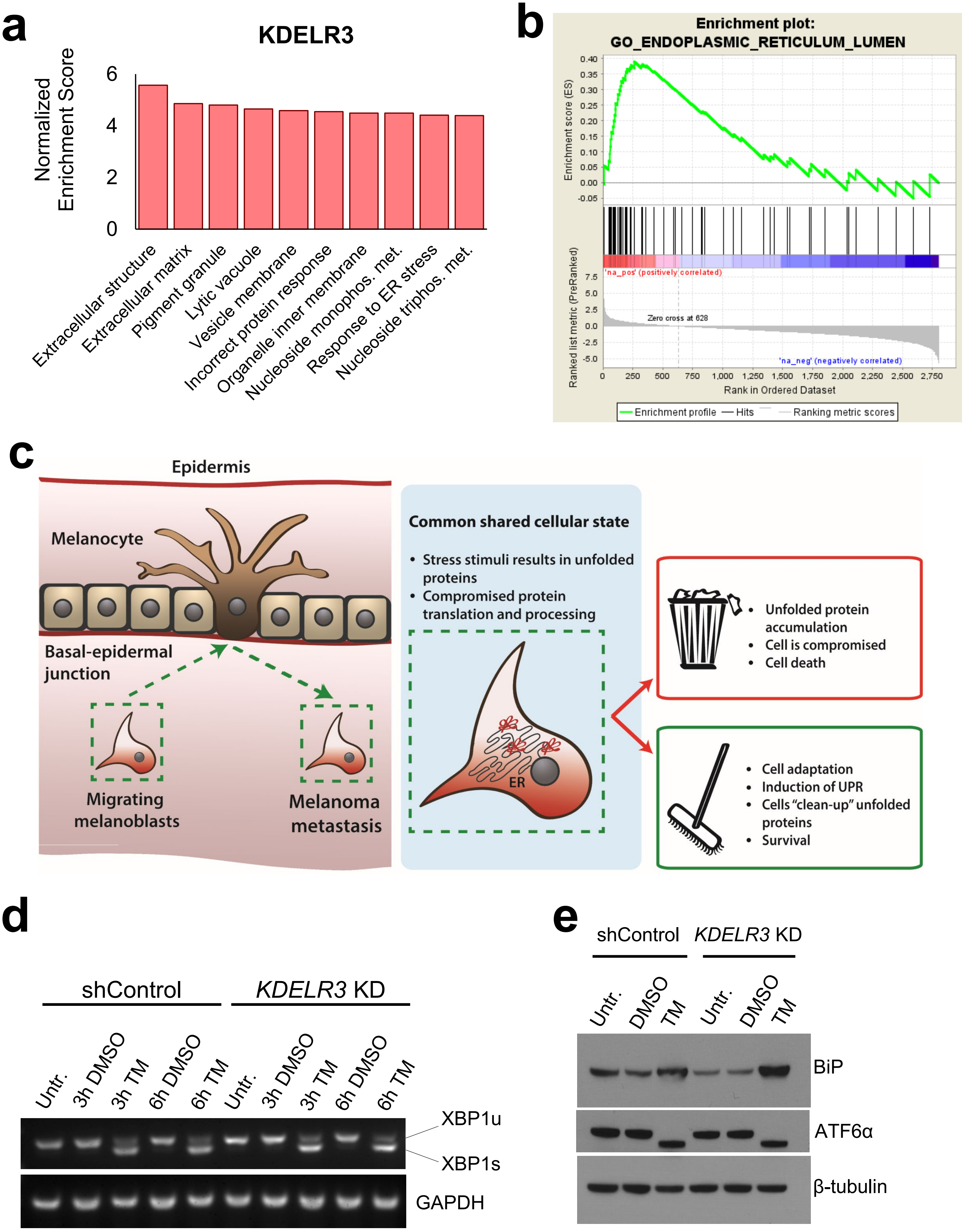
a
KDELR3
Normalized Enrichment Score
0
2
4
6
Extracellular structure
Extracellular matrix
Pigment granule
Lytic vacuole
Vesicle membrane
Incorrect protein response
Organelle inner membrane
Nucleoside monophos. met.
Response to ER stress
Nucleoside triphos. met.
b
Enrichment plot:
GO_ENDOPLASMIC_RETICULUM_LUMEN
Enrichment score (ES)
Ranked list metric (PreRanked)
'na_pos' (positively correlated)
Zero cross at 628
'na_neg' (negatively correlated)
Rank in Ordered Dataset
Enrichment profile
Hits
Ranking metric scores
c
Epidermis
Melanocyte
Basal-epidermal junction
Migrating melanoblasts
Melanoma metastasis
Common shared cellular state
Stress stimuli results in unfolded proteins
Compromised protein translation and processing
ER
Unfolded protein accumulation
Cell is compromised
Cell death
Cell adaptation
Induction of UPR
Cells "clean-up" unfolded proteins
Survival
d
shControl
KDELR3 KD
Untr.
3h DMSO
3h TM
6h DMSO
6h TM
XBP1u
XBP1s
GAPDH
e
shControl
KDELR3 KD
Untr.
DMSO
TM
BiP
ATF6α
β-tubulin

**Supplementary Figure 6: *KDELR3* and the ER Stress Response in metastatic melanoma. a**, GSEA of gene co-expression within skin cutaneous melanoma patients of the TCGA (n = 479). Top 10 *KDELR3*-associated GO pathways represented, FDR < 0.0001. Go pathways in order: Extracellular structure organization; Extracellular matrix; Pigment granule; Lytic vacuole; Vesicle membrane; Response to topologically incorrect protein; Organelle inner membrane; Nucleoside monophosphate metabolic process; Response to endoplasmic reticulum stress; Nucleoside triphosphate metabolic process. **b**, Gene Ontology (GO) term enrichment of positively enriched pathways following proteomic analysis of si*KDELR3* knockdown cells (1205Lu) compared to parental cells and non-targeting control cells identified GO_ENDOPLASMIC_RETICULUM_LUMEN as the GeneSet consistently enriched in two independent replicates. Enrichment Score (ES) = 0.39025578, Normalized Enrichment Score (NES) = 1.823677, Nominal p-value < 0.0001, FDR q-value = 0.09752231, FWER p-value = 0.186. Representative results from two independent experiments. **c**, Migrating melanoblasts and metastasizing melanoma cells are affected by similar stress stimuli[34], i.e. low oxygen/nutrient supply, foreign microenvironment, detachment, shear stressors. Sub-optimal protein translation and processing during times of cellular stress results in build-up of unfolded proteins in the ER-Golgi transport network (ER stress). Unfolded protein accumulation further compromises normal protein translation, processing and transport, thereby affecting cellular function. Inability to cope with ER stress typically results in cell death. Induction of the unfolded protein response (UPR) is a cell survival mechanism that allows cells to adapt to ER stress[77]. **d**, RT-PCR analysis of XBP1 splicing in shRNA non-targeting control and sh*KDELR3* knockdown (1205Lu cells). Un-spliced, inactive XBP1 (XBP1u) was compared to spliced, active XBP1 signaling (XBP1s) Cells were treated for the time specified with: untreated, DMSO, or 3µg/ml Tunicamycin (TM). GAPDH loading control. Representative results from two independent experiments in two different cell lines (WM-46 and 1205Lu)**. e**, BiP and ATF6α protein expression in non-targeting control (shControl) and sh*KDELR3* (*KDELR3* KD) WM-46 cells in untreated (Untr.), DMSO controls (DMSO) and treated with 2.5 µg/ml Tunicamycin (TM) 15 hours before collection. BiP: representative results from three independent cell lines (1205Lu, WM-46, B16). ATF6α: representative results from three independent experiments in two cell lines (WM-46 and 1205Lu).

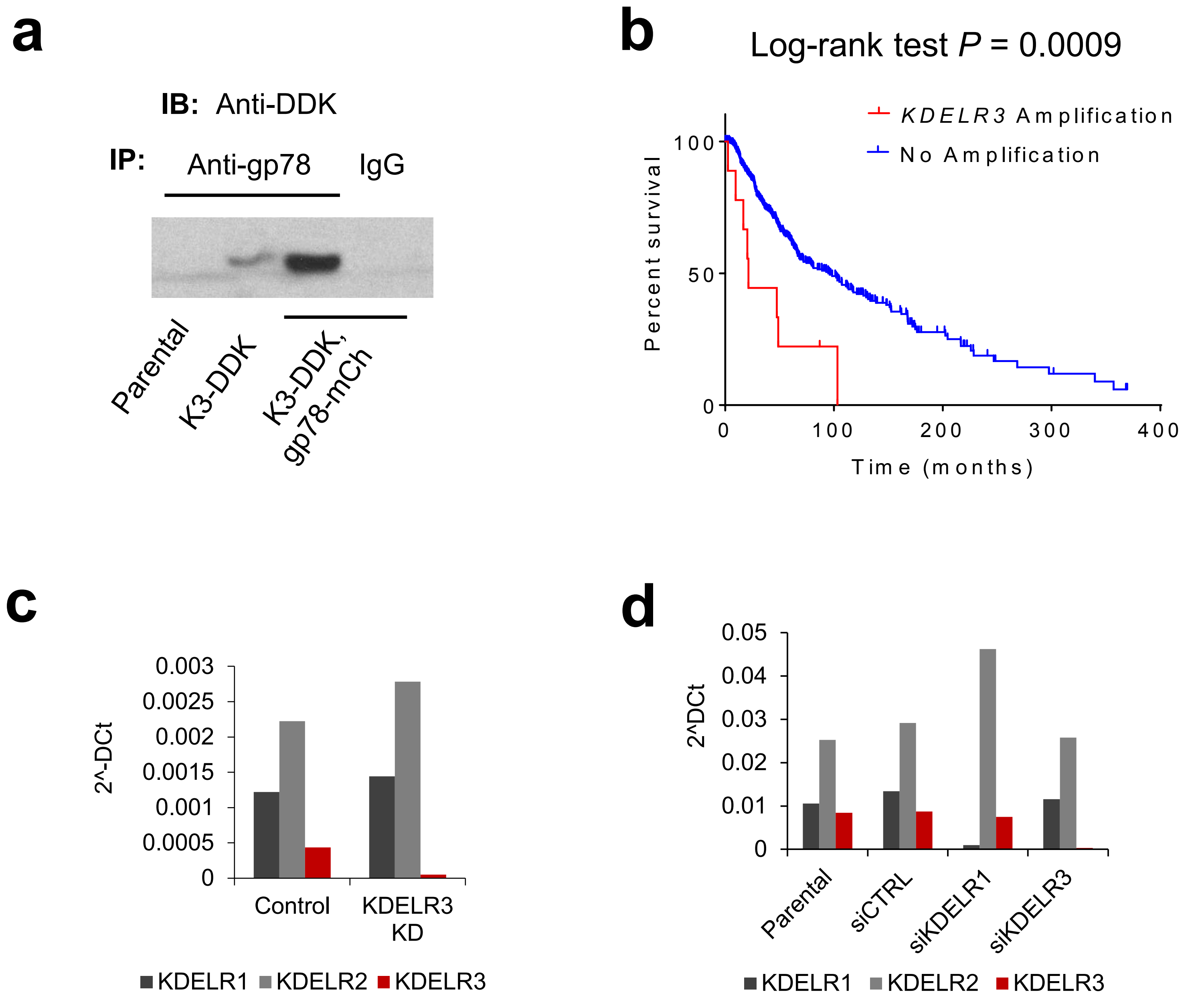


**Supplementary Figure 7: KDELR3 biology in melanoma. a**, Co-immunoprecipitation of endogenous gp78 and mCherry tagged gp78 (gp78-mCh) with FLAG-tagged KDELR3 (K3-DDK) in stably transduced 1205Lu cells. **b**, Kaplan-Meier curve of skin cutaneous melanoma patient (TCGA-SKCM) overall survival with (N=9) and without (N=349) *KDELR3* CNV amplifications; CNV determined by GISTIC 2.0. Data generated by cBioPortal for Cancer Genomics. Log-rank test, $P$ = 0.009. **c-d**, KDELR family member expression in 1205Lu metastatic melanoma cells following shRNA (**c**) or siRNA knockdown (**d**; day 3 post knockdown). **a**, **c-d**, Representative of three independent experiments.

| Gene comparison | R-squared | 1/slope | P-value | Significance ? |
|---|---|---|---|---|
| **Kdelr3 vs P4ha2** | 0.5902 | 0.4608 | < 0.0001 | Yes |
| **Kdelr3 vs Gulp1** | 0.02636 | -5.53 | 0.3169 | No |
| **Kdelr3 vs Dab2** | 0.4415 | 0.6481 | < 0.0001 | Yes |
| **P4ha2 vs Gulp1** | 0.02643 | 15.6 | 0.3162 | No |
| **P4ha2 vs Dab2** | 0.3827 | 1.966 | < 0.0001 | Yes |
| **Gulp1 vs Dab2** | 0.00099 | -15.26 | 0.8473 | No |

**Supplementary Table 1: Co-expression of 4 functionally validated genes within four independent mouse melanoma models.** Red text denotes significant positive correlation.

| Gene Comparison | Spearman's Correlation | p-Value | q-Value | Significance? |
|---|---|---|---|---|
| KDELR3 vs P4HA2 | 0.435 | 3.49e-23 | 4.40e-20 | Yes |
| KDELR3 vs DAB2 | 0.213 | 2.999e-6 | 4.169e-5 | Yes |
| KDELR3 vs GULP1 | -0.0201 | 0.662 | 0.774 | No |
| P4HA2 vs GULP1 | -0.00202 | 0.965 | 0.978 | No |
| P4HA2 vs DAB2 | 0.173 | 1.614e-4 | 8.925e-4 | Yes |
| GULP1 vs DAB2 | -0.273 | 0.1.74e-9 | 8.39e-9 | Yes |

**Supplementary Table 2: Co-expression of 4 functionally validated genes within melanoma patients.** TCGA, cBioPortal. Red text denotes significant positive correlation. Blue text denotes significant negative correlation.

| NAME | SIZE | ES | NES | NOM p-val | FDR q-val | FWER p-val | RANK AT MAX |
|---|---|---|---|---|---|---|---|
| **GO_ENDOPLASMIC_RETICULUM** | 364 | 0.205073 | 4.232328 | 0 | 0 | 0 | 1219 |
| **GO_ENDOPLASMIC_RETICULUM_PART** | 282 | 0.22867 | 4.218896 | 0 | 0 | 0 | 1122 |
| **GO_ENDOPLASMIC_RETICULUM_LUMEN** | 58 | 0.466685 | 4.165108 | 0 | 0 | 0 | 1094 |
| **GO_NUCLEAR_OUTER_MEMBRANE_ENDOPLASMIC_RETICULUM_MEMBRANE_NETWORK** | 242 | 0.178756 | 3.147752 | 0 | 0 | 0 | 1122 |
| **GO_EXTRACELLULAR_STRUCTURE_ORGANIZATION** | 56 | 0.330455 | 2.896429 | 0 | 6.30E-04 | 0.004 | 1006 |
| GO_VESICLE_COATING | 37 | 0.399024 | 2.860223 | 0 | 7.80E-04 | 0.006 | 1053 |
| **GO_GOLGI_APPARATUS_PART** | 155 | 0.188323 | 2.639097 | 0 | 0.006592 | 0.058 | 1053 |
| **GO_GOLGI_MEMBRANE** | 133 | 0.201222 | 2.582831 | 0.001934 | 0.008083 | 0.081 | 1046 |
| GO_SINGLE_ORGANISM_MEMBRANE_BUDDING | 37 | 0.346847 | 2.525803 | 0 | 0.009166 | 0.112 | 1046 |
| **GO_GOLGI_APPARATUS** | 254 | 0.142598 | 2.545466 | 0 | 0.009326 | 0.105 | 1075 |
| GO_TRANSPORT_VESICLE | 73 | 0.249239 | 2.49897 | 0 | 0.010362 | 0.136 | 1046 |
| GO_VESICLE_TARGETING | 38 | 0.3275 | 2.400361 | 0 | 0.017508 | 0.245 | 1046 |
| **GO_ENDOPLASMIC_RETICULUM_GOLGI_INTERMEDIATE_COMPARTMENT** | 41 | 0.321227 | 2.40303 | 0 | 0.018265 | 0.236 | 1055 |
| GO_MEMBRANE_BUDDING | 49 | 0.288477 | 2.385064 | 0 | 0.018397 | 0.278 | 1053 |
| GO_PROTEINACEOUS_EXTRACELLULAR_MATRIX | 39 | 0.321062 | 2.357299 | 0 | 0.021642 | 0.339 | 945 |
| GO_LOCALIZATION_WITHIN_MEMBRANE | 43 | 0.301722 | 2.322892 | 0.001916 | 0.025717 | 0.417 | 893 |
| GO_MITOCHONDRIAL_PART | 312 | 0.116061 | 2.274941 | 0 | 0.033188 | 0.523 | 1280 |
| GO_ER_TO_GOLGI_TRANSPORT_VESICLE | 25 | 0.376966 | 2.246624 | 0 | 0.036413 | 0.594 | 1043 |
| GO_TRANSPORT_VESICLE_MEMBRANE | 27 | 0.359304 | 2.249874 | 0 | 0.037419 | 0.583 | 1139 |
| GO_GLYCOPROTEIN_METABOLIC_PROCESS | 51 | 0.259467 | 2.224952 | 0.002041 | 0.040386 | 0.637 | 1218 |

**Supplementary Table 3: Gene Set Enrichment Analysis (GSEA) of positively enriched pathways following si*KDELR3* knockdown compared to non-targeting control cells.** Gene Ontology (GO) term enrichment. FDR < 0.05 denotes significant pathways. ES, Enrichment Score; NES, Normalized Enrichment Score. Bold font denotes pathways consistent with siAMFR knockdown.

| NAME | SIZE | ES | NES | NOM p-val | FDR q-val | FWER p-val | RANK AT MAX |
|---|---|---|---|---|---|---|---|
| **GO_ENDOPLASMIC_RETICULUM** | 364 | 0.191889 | 4.097375 | 0 | 0 | 0 | 1465 |
| **GO_ENDOPLASMIC_RETICULUM_PART** | 282 | 0.213989 | 3.958402 | 0 | 0 | 0 | 1033 |
| **GO_NUCLEAR_OUTER_MEMBRANE_ENDOPLASMIC_RETICULUM_MEMBRANE_NETWORK** | 242 | 0.19813 | 3.406401 | 0 | 0 | 0 | 1465 |
| **GO_ENDOPLASMIC_RETICULUM_LUMEN** | 58 | 0.309021 | 2.7944 | 0 | 0.002871 | 0.013 | 1371 |
| GO_PROTEIN_LOCALIZATION_TO_ENDOPLASMIC_RETICULUM | 93 | 0.222677 | 2.561967 | 0 | 0.009951 | 0.067 | 1945 |
| **GO_ENDOPLASMIC_RETICULUM_GOLGI_INTERMEDIATE_COMPARTMENT** | 41 | 0.342655 | 2.567371 | 0 | 0.01106 | 0.062 | 998 |
| **GO_EXTRACELLULAR_STRUCTURE_ORGANIZATION** | 56 | 0.283743 | 2.482635 | 0 | 0.018637 | 0.143 | 599 |
| GO_PROTEIN_N_LINKED_GLYCOSYLATION | 21 | 0.432285 | 2.323848 | 0 | 0.030658 | 0.406 | 1027 |
| **GO_GOLGI_APPARATUS** | 254 | 0.131271 | 2.32819 | 0 | 0.032158 | 0.404 | 1443 |
| GO_MULTI_ORGANISM_METABOLIC_PROCESS | 103 | 0.206993 | 2.401925 | 0.001996 | 0.03278 | 0.265 | 1900 |
| GO_VESICLE_COAT | 24 | 0.395935 | 2.297542 | 0 | 0.03318 | 0.475 | 1415 |
| GO_GOLGI_VESICLE_TRANSPORT | 111 | 0.196268 | 2.332478 | 0 | 0.033759 | 0.394 | 1486 |
| GO_REGULATION_OF_ENDOTHELIAL_CELL_PROLIFERATION | 15 | 0.504046 | 2.368323 | 0 | 0.034191 | 0.323 | 986 |
| GO_PROTEIN_TARGETING_TO_MEMBRANE | 97 | 0.198248 | 2.301479 | 0 | 0.034262 | 0.464 | 1961 |
| **GO_GOLGI_APPARATUS_PART** | 155 | 0.160576 | 2.338985 | 0.002004 | 0.035189 | 0.385 | 688 |
| GO_ESTABLISHMENT_OF_PROTEIN_LOCALIZATION_TO_ENDOPLASMIC_RETICULUM | 87 | 0.217569 | 2.348091 | 0 | 0.036006 | 0.368 | 1945 |
| GO_POSITIVE_REGULATION_OF_EPITHELIAL_CELL_PROLIFERATION | 16 | 0.486615 | 2.369748 | 0 | 0.037693 | 0.32 | 887 |
| **GO_GOLGI_MEMBRANE** | 133 | 0.170127 | 2.230692 | 0 | 0.048454 | 0.636 | 1004 |

**Supplementary Table 4: Gene Set Enrichment Analysis (GSEA) of positively enriched pathways following si*AMFR* knockdown compared to non-targeting control cells.** Gene Ontology (GO) term enrichment. FDR < 0.05 denotes significant pathways. ES, Enrichment Score; NES, Normalized Enrichment Score. Bold font denotes pathways consistent with si*KDELR3* knockdown.
